# STXBP5/tomosyn regulates the small RhoA GTPase to control the dendritic stability of neurons and the surface expression of AMPA receptors

**DOI:** 10.1101/617845

**Authors:** Wenjuan Shen, Michaela B.C. Kilander, Morgan S. Bridi, Jeannine A. Frei, Robert F. Niescier, Shiyong Huang, Yu-Chih Lin

## Abstract

Tomosyn, a protein encoded by syntaxin-1-binding protein 5 (*STXBP5*) gene, has a well-established presynaptic role in the inhibition of neurotransmitter release and the reduction of synaptic transmission by its conical interaction with the soluble N-ethylmaleimide-sensitive factor attachment protein receptor (SNARE) machinery. The postsynaptic role of tomosyn in dendritic arborization, spine stability, and trafficking of ionotropic glutamate receptors remains to be elucidated. We used short hairpin RNA (shRNA) to knock down tomosyn in mouse primary neurons to evaluate the postsynaptic cellular function and molecular signaling regulated by tomosyn. Knockdown of tomosyn led to an increase of RhoA GTPase activity accompanied by compromised dendritic arborization, loss of dendritic spines, decreased surface expression of AMPA receptors, and reduced miniature excitatory postsynaptic current (mEPSC) frequency. Inhibiting RhoA signaling was sufficient to rescue the abnormal dendritic morphology and the surface expression of AMPA receptors. The function of tomosyn regulating RhoA is mediated through the N-terminal WD40 motif, where two variants each carrying a single nucleotide mutation in this region, were found in individuals with autism spectrum disorder (ASD). We demonstrated that these variants displayed loss-of-function phenotypes. Unlike the wild-type tomosyn, these two variants failed to restore the reduced dendritic complexity, spine density, as well as decreased surface expression of AMPA receptors in tomosyn knockdown neurons. This study uncovers a critical role of tomosyn, independent of its interaction with the SNARE machinery, in maintaining neuronal function by inhibiting RhoA activity. Further analysis of tomosyn variants also provides a potential mechanism for explaining cellular pathology in ASD.

**Significance Statement:** This study unveils a vital role of tomosyn in the maintenance of neuronal morphology, basal synaptic transmission, and AMPA receptor surface expression that is distinct from its presynaptic role. Tomosyn affects dendritic stability and glutamate receptor trafficking via the regulation of the Rho signaling pathway and this interaction is likely independent of the interaction with the dendritic SNARE complex, such as syntaxin-4. The WD40 domain of tomosyn is necessary to conduct the Rho regulation and two autism-associated variants localized at the WD40 domain perturb this function. The current study reveals a novel molecular link between dendritic stability and synaptic function, which could advance a greater understanding of the cellular pathologies involved in neurodevelopmental disorders, such as ASD.

## Introduction

Formation and maintenance of neuronal connections are crucial for promoting proper brain function. Failure of these processes are often observed in neurological disorders accompanied by detrimental behaviors. Excitatory neurons are major projection cells in the brain and extend complex dendritic arbors to establish extensive connections with other neurons. Mature neurons have actin-rich spines that protrude from dendrites and are the major structures harboring synaptic components. Cytoskeleton rearrangement by Rho-signaling pathways is the driving force that regulates neuronal stability and synaptic plasticity (*1–3*). Ionotropic glutamate receptors, including AMPA and NMDA receptors, cluster primarily on dendritic spines, and are responsible for excitatory synaptic transmission and synaptic plasticity (*4–6*). Understanding the mechanistic regulations between the cytoskeletal machinery and glutamate receptors trafficking will allow us to better understand the underlying mechanisms that promote neuronal stability and function.

Tomosyn was first identified as a protein encoded by syntaxin-1-binding protein 5 (STXBP5) in rats (*7*). Structurally, tomosyn contains an N-terminal WD40-repeats domain and a C-terminal coiled-coil motif (*8*). The R-SNARE-like structure at the C-terminus allows tomosyn to inhibit the formation of the SNARE complex by competing with Munc18 for binding to syntaxin-1, thereby negatively regulating neurotransmitter release (*7, 9, 10*). Mice overexpressing tomosyn in the hippocampus showed impaired spatial learning and memory, potentially via reduced synaptic transmission in mossy fiber (MF)-CA3 synapses (*11*). Conversely, tomosyn knockout mice exhibited accelerated kindling behaviors due to increased glutamate release in the hippocampal dentate gyrus (*12*). Selectively knocking down tomosyn in MF-CA3 synapses also impaired facilitation, long-term potentiation (LTP), and PKA-induced potentiation (*9*). Despite accumulating evidence for the presynaptic roles of tomosyn, relatively few studies have focused on its role in postsynaptic compartments. One study has shown that tomosyn regulates neurite outgrowth in immature neurons by strongly binding to Rho-associated serine/threonine kinase (ROCK)-phosphorylated syntaxin-1 (*13*). Tomosyn has also been shown to regulate dendritic morphology via the ubiquitin-proteasome system (*14*). Here, we provide evidence showing that tomosyn regulates RhoA signaling and the trafficking of glutamate receptors to maintain the structural stability of neurons.

Mechanisms that regulate neuronal stability and synaptic function are often vulnerable targets in neurodevelopmental disorders, such as ASD (*15–17*). Individuals with ASD commonly exhibit difficulties in social communication and interaction, repetitive behaviors, and restricted interests (*18*). Although ASD exhibits a high degree of genetic heterogeneity, many identified risk genes converge on similar cellular pathways, including those that regulate neurite outgrowth, spine stability, synaptic plasticity, excitatory/inhibitory balance, and trafficking of glutamate receptors (*19–27*). Several human genetic studies, including genome-wide association studies and whole-exome sequencing, have linked the deletion, as well as two tomosyn variants (L412V and Y502C) to ASD (*28–31*). In the present study, reductions of dendritic arborization and spine density accompanied by enhanced RhoA activity and decreased surface expression of glutamate receptors were observed in tomosyn knockdown neurons. ASD-associated tomosyn L412V and Y502C variants displayed a loss-of-function effect and failed to restore the cellular defects in tomosyn knockdown neurons. Our study reveals a pivotal postsynaptic function of tomosyn, which mediates a potential pathway for maintaining the structural stability of neurons, and may be altered in ASD.

## Results

### Tomosyn is expressed at pre- and post-synaptic areas

Protein expression of tomosyn in developing mouse brains was first determined using Western blot analysis. The expression of tomosyn in the hippocampus was readily detectable from P0 and increased temporally during development (Fig. 1A). Tomosyn levels reached a four-fold elevation by P14, suggesting an increasing role during neuronal maturation. Spatial analysis at P14 revealed a dominant expression profile of tomosyn in the forebrain compared to the cerebellum (fig. S1A). Further examination in whole brain also showed that the protein expression level of tomosyn increased nearly two-fold at P14 compared to P0 (fig. S1B). In cultured cortical neurons, tomosyn also showed a gradual increase of expression with PSD-95, although not to the same degree, suggesting a role in synaptogenesis (fig. S1C). To determine the subcellular localization of tomosyn, synaptic fractionation of mouse forebrain was first performed (Fig. 1B). Tomosyn was enriched in presynaptic compartments including syntaxin-1-rich synaptic plasma membrane (SPM) and synaptic vesicles (S3). However, tomosyn was also detected in the postsynaptic density (PSD), indicated by the strong presence of PSD-95. In addition, cultured hippocampal neurons were co-immunostained for tomosyn and the dendritic marker, microtubule-associated protein 2 (MAP2). Endogenous tomosyn was shown to localize in both the soma and MAP2-positive dendrites (Fig. 1C). Overexpressed tomosyn-GFP also localized to dendritic compartments, including dendritic spines (Fig. 1D). The expression level of tomosyn in dendritic spines was similar to dendrite shafts (Fig. 1E). Furthermore, immunofluorescence labeling of cultured hippocampal neurons revealed tomosyn expression in both CaMKIIα-positive excitatory neurons and GAD67-positive inhibitory neurons (fig. S1D). These data establish the spatiotemporal expression pattern of tomosyn in the brain and suggest a potential postsynaptic role in dendritic development.

**Fig. 1.**
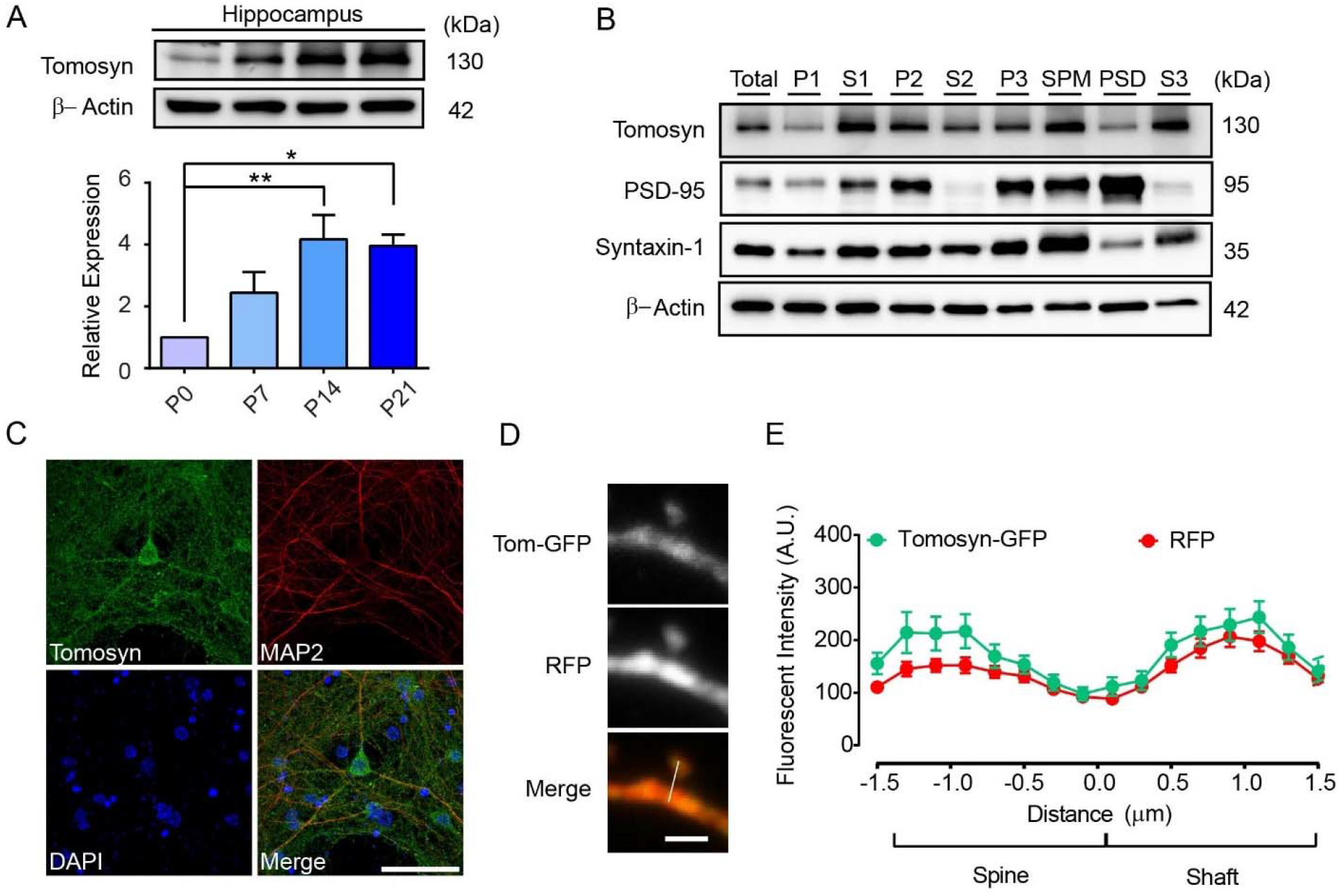
Tomosyn is localized in both pre- and post-synaptic compartments. (**A**) Western blot analysis shows the developmental expression pattern of tomosyn protein in the hippocampus at P0, P7, P14, and P21. (**B**) Synaptic fractionation of forebrain lysates (total) shows the subcellular localization of tomosyn in P21 mouse brains. P1, nuclei; S1, cytosol/membranes; P2, crude synaptosomes; S2, cytosol/light membranes; P3, synaptosomes; SPM, synaptic plasma membranes; PSD, postsynaptic density; S3, synaptic vesicles. (**C**) Confocal immunofluorescent images showing tomosyn (green), MAP2 (red), and DAPI (blue) from a cultured hippocampal neuron at 15 DIV. Scale bar, 50 μm. (**D**) Confocal images of a dendritic spine of a cultured neuron transfected with RFP and tomosyn-GFP. (**E**) Fluorescence intensity of tomosyn-GFP and RFP in the dendritic spine versus the dendritic shaft was quantified by line scan (shown in **D**). Three independent experiments are conducted in **A-B**. 29 spines are analyzed in **E**.

### Knockdown of tomosyn decreases dendritic complexity and dendritic spine density

To examine the effect of tomosyn on neuronal morphology, a short hairpin RNA (shRNA) expression system was used to knock down tomosyn in cultured hippocampal neurons. The knockdown efficiency of two shRNAs targeting tomosyn, shRNA482 and shRNA1083, was first determined in N2a cells. shRNA482 and shRNA1083 resulted in 87.9% and 79.3% reduction, respectively, in the amount of overexpressing GFP-tagged tomosyn protein compared to scrambled shRNA (fig. S2A). shRNA482, which most effectively decreased tomosyn expression, was chosen for all subsequent experiments and is referred to as shTomosyn hereafter. To further evaluate the knockdown specificity of shTomosyn, an shRNA-resistant form of tomosyn was engineered as Tom^r^-GFP. Co-expressing Tom^r^-GFP with shTomosyn effectively restored the protein level of tomosyn back to control levels (fig. S2B). shTomosyn was then used to knock down tomosyn expression in cultured hippocampal neurons, and their dendritic arborization was examined. Reconstructions of dendritic trees (Fig. 2A) and Sholl analysis (Fig. 2B) showed that the overall size of the dendritic arbors was smaller in tomosyn knockdown neurons. Co-expression of Tom^r^-GFP efficiently restored dendritic complexity within the first 100 µm radius of dendritic arbors in knockdown neurons. Knockdown of tomosyn also resulted in decreased spine density (scramble: 0.61 ± 0.02 /µm; shTomosyn: 0.36 ± 0.02 /µm), while re-expressing Tom^r^-GFP (0.53 ± 0.02 /µm) successfully rescued dendritic spine loss (Fig. 2C and D). Together, these data suggest that tomosyn is required for dendritic arborization and dendritic spine formation.

**Fig. 2.**
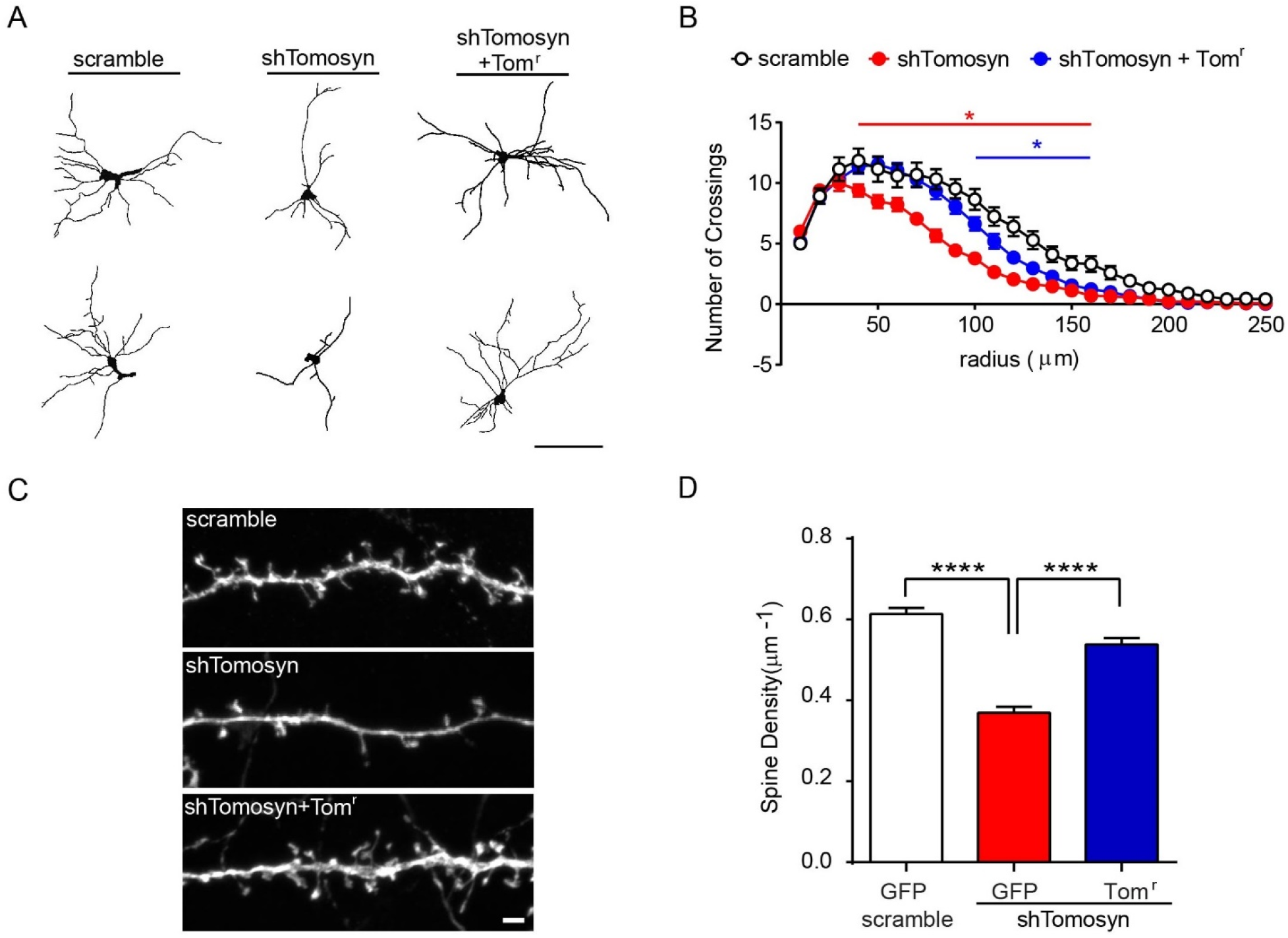
Knockdown of tomosyn reduces dendritic complexity and dendritic spine density. (**A**) Representative images of reconstructed dendritic trees from mouse hippocampal neurons transfected with scrambled shRNA + GFP or shTomosyn + GFP or shTomosyn + Tom^r^-GFP. Scale bar, 100 µm. (**B**) Sholl analysis of reconstructed neurons. Neurons expressing shTomosyn exhibited compromised dendritic complexity, whereas expression of Tom-GFP rescued simplified dendritic complexity in shTomosyn-expressing neurons compared to scramble controls. *indicates significant difference at a particular radius compared to scramble controls, two-way ANOVA with Tukey’s test. n =23 ∼ 25 neurons from three different cultures. (**C**) Confocal fluorochrome images of dendritic spines from neurons transfected with scrambled shRNA + GFP, shTomosyn + GFP, or shTomosyn + Tom^r^-GFP for 72hrs. Scale bar, 2 µm. (**D**) Mean spine density was decreased in tomosyn knockdown neurons, whereas spine density was restored in Tom^r^-GFP and shTomosyn-co-expressing neurons compared to shTomosyn + GFP. \*\*\*\**p* < 0.0001 by one-way ANOVA with Dunnett’s multiple comparisons test. n = 29 ∼ 31 neurons from three different cultures.

### Tomosyn deficiency leads to reduced frequency of mEPSCs

Because tomosyn deficiency results in abnormal dendritic arborization and spinogenesis, we asked whether tomosyn plays a role in maintaining basal excitatory synaptic transmission. To answer this question, the effect of tomosyn knockdown on AMPA receptor (AMPAR)-mediated miniature excitatory postsynaptic currents (mEPSCs) in cultured hippocampal neurons was examined. mEPSC frequency was significantly reduced in tomosyn knockdown cells compared to controls (Fig. 3C, scramble: 5.18 ± 0.78 Hz, shTomosyn: 1.46 ± 0.36 Hz). Charge (Fig. 3E, scramble: 42.41 ± 3.61, shTomosyn: 32.89 ± 2.23), but not amplitude (Fig. 3D, scramble: 16.35 ± 1.46 pA, shTomosyn: 14.55 ± 1.01 pA), was decreased in tomosyn knockdown neurons compared to scrambled shRNA transfected neurons. Analysis of mEPSC kinetics revealed that the decay time constant was significantly smaller in tomosyn knockdown neurons than in the scramble controls (Fig. 3F, scramble: 3.7 ± 0.25 ms, shTomosyn: 2.8 ± 0.21 ms). These data showing a reduction of mEPSC frequency support the spine loss phenotype observed in tomosyn knockdown neurons, while the attenuation in the decay time constant may indicate a change in AMPAR subunit composition at the synapse in shTomosyn neurons.

**Fig. 3.**
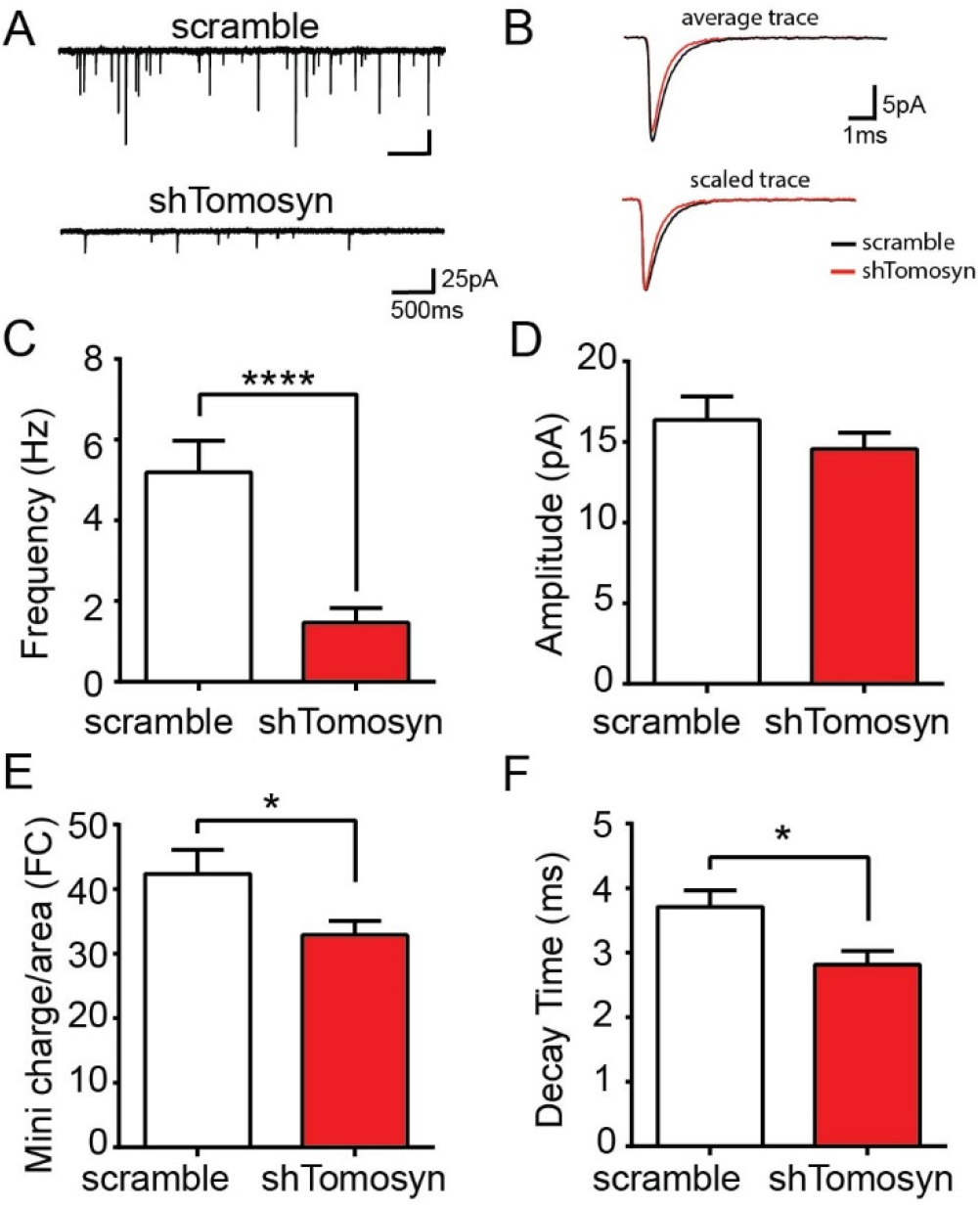
Frequency of miniature EPSCs is decreased in tomosyn knockdown neurons. Cultured hippocampal neurons were transfected with either scrambled shRNA or shTomosyn. (**A**) An example of current traces collected from whole-cell voltage-clamp of hippocampal neurons. (**B**) Examples of average (top) and scaled (bottom) mEPSC traces from scrambled shRNA and shTomosyn transfected neurons. Summary graphs show (**C**) frequency (\*\*\*\**p* < 0.0001) (**D**) amplitude (*p* = 0.32), and (**E**) charge (\**p* < 0.05) of recorded mEPSCs. Summary graph showing kinetic properties of mEPSCs, including (**F**) decay time constant (\**p* < 0.05). Statistical analysis was conducted by Mann-Whitney test for **C** and by unpaired student’s *t* test with Welch’s correction for **D-F**. n = 22 neurons for scramble, n = 21 neurons for shTomosyn from five independent cultures.

### Tomosyn knockdown leads to elevated RhoA GTPase activity

Decreased dendritic arborization and destabilization of dendritic spines on pyramidal neurons have previously been linked to the increased activity of RhoA GTPase (*2*). Thus, we investigated whether the simplification of dendritic arbors and loss of dendritic spines observed in neurons with tomosyn knockdown were the result of elevated RhoA activity. Hippocampal neurons were co-transfected with either scrambled shRNA or shTomosyn, as well as an intramolecular Förster Resonance Energy Transfer (FRET) RhoA biosensor, which contains the CFP (donor)/YFP (acceptor) FRET pair flanked by full-length RhoA on one side, and the Rho-binding domain of the effector rhotekin (RBD) on the other side (*32*). Upon activation of RhoA, the RBD element associates with RhoA-GTP, thereby causing the biosensor molecule to change its conformation into a FRET-favorable state. Hence, RhoA activity can be monitored by measuring the FRET occurring between the CFP and YFP fluorophores (*32, 33*). FRET efficiency was recorded by two well-established techniques: sensitized emission (Fig. 4A and B) and acceptor photobleaching (fig. S3A-C). Whole-cell RhoA biosensor FRET signal recorded by sensitized emission revealed that knockdown of tomosyn resulted in a significantly higher FRET ratio compared to scramble controls (Fig. 4A and B). Similarly, acceptor (YFP) photobleaching FRET (pbFRET) performed at selected regions along apical and basal dendrites showed that overall E_FRET_ was greater in tomosyn knockdown neurons compared to scramble controls (fig. S3A-C). However, within each sample group (scramble and shTomosyn) no significant differences in E_FRET_ were observed between the specified dendritic compartments (fig. S3C). Collectively, these data show that RhoA activity is increased in tomosyn knockdown neurons, indicating that tomosyn might play a role in the maintenance of dendritic morphology by inhibiting RhoA GTPase signaling.

**Fig. 4.**
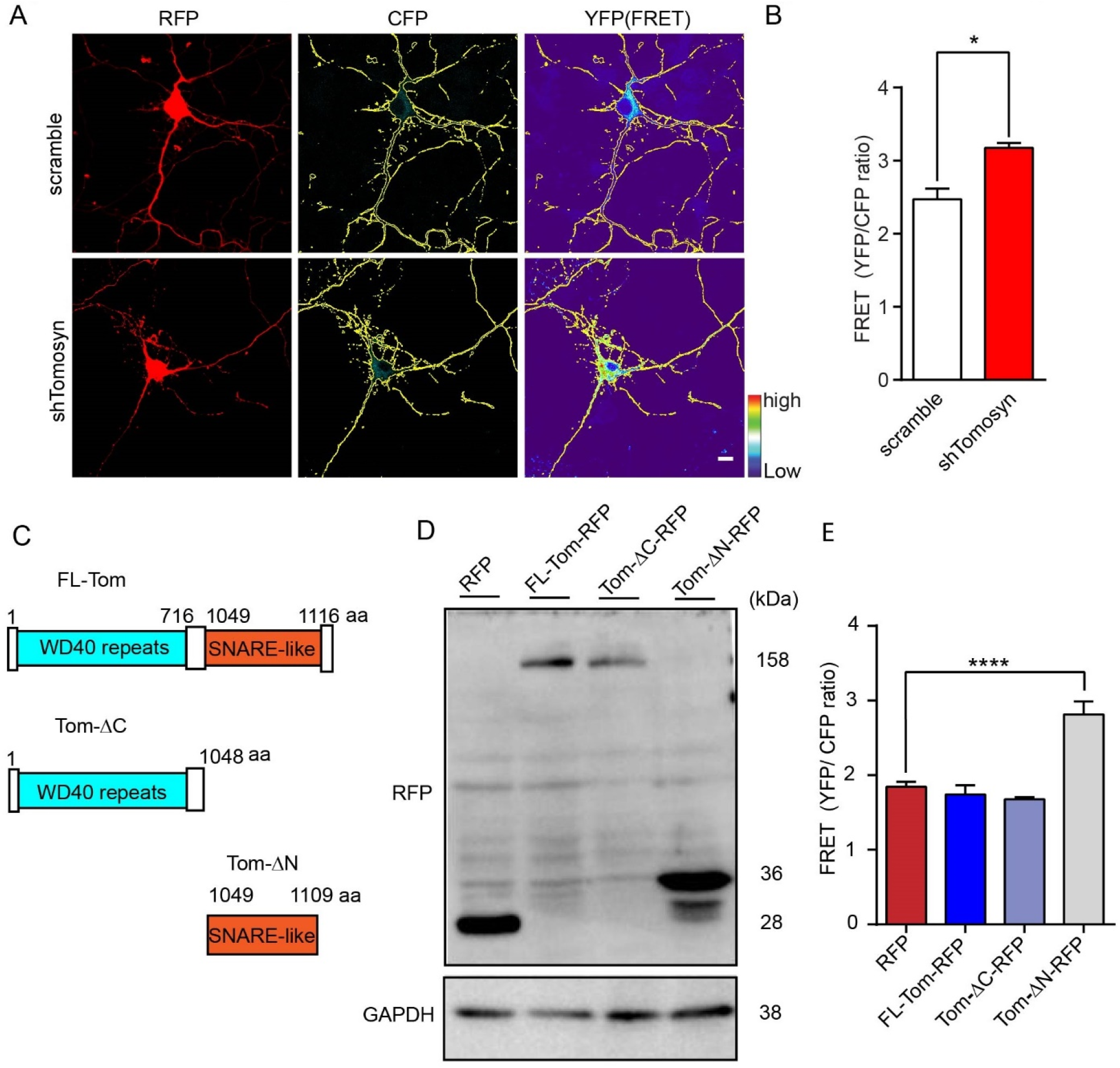
Tomosyn knockdown results in increased RhoA activity in hippocampal neurons. Hippocampal neurons were co-transfected with a RhoA Biosensor and either scrambled shRNA or shTomosyn. (**A**) Representative images show the neurons in the intensity-modulated display mode. Scale bar, 10 µm. (**B**) FRET efficiency was determined as a ratio of YFP/CFP. n = 69 ∼ 72 neurons per condition from four independent experiments. \**p* < 0.05 by two-tailed Mann-Whitney test. (**C**) Schematic structure of domain mutant tomosyn. FL-Tom, full-length mouse tomosyn (1-1166 aa); Tom-ΔC, containing an N-terminal domain with WD40 repeats (1-1048 aa); Tom-ΔN, containing a SNARE coil-coiled domain (1049-1109 aa). (**D**) Western blot shows expression of domain mutant tomosyn (Tom-ΔC-RFP and Tom-ΔN-RFP), FL-Tom-RFP and RFP control in N2a cells. (**E**) Representative graph showing FRET efficiency in domain mutant tomosyn-expressing neurons. Neurons with Tom-ΔN-RFP showed increased FRET efficiency. \*\*\*\**p* < 0.0001 compared to RFP vector by one-way ANOVA with Dunnett’s multiple comparisons test. n = 28 ∼ 32 neurons per condition from four independent experiments.

To further investigate which part of tomosyn is essential for inhibiting RhoA activity, two domain mutants were constructed: Tom-ΔC-RFP, which contains the N-terminal WD40 domain, and Tom-ΔN-RFP, which contains only the R-SNARE domain of tomosyn (Fig. 4C). Western blot analysis showed that both Tom-ΔC-RFP and Tom-ΔN-RFP migrated as shorter versions of tomosyn at the predicted size (Fig. 4D). FRET analysis showed that RhoA activity in neurons expressing Tom-ΔC-RFP is comparable to that of RFP control and full-length tomosyn. However, E_FRET_ was significantly increased in neurons expressing Tom-ΔN-RFP (Fig. 4E), suggesting the lack of the WD40 domain of tomosyn is sufficient to increase RhoA activity. Taken together, these data suggest that tomosyn likely functions as a negative regulator for RhoA activity via the WD40 domain in neurons.

### Inhibition of RhoA signaling restores dendritic arborization and spine density in tomosyn-deficient neurons

To determine whether inhibiting RhoA activity is sufficient to rescue the aberrant dendritic phenotypes in tomosyn knockdown neurons, dominant-negative RhoA (T19N-RhoA) was co-transfected in scramble control- and shTomosyn-expressing neurons (Fig. 5A and B). In agreement with previous findings, WT-RhoA or T19N-RhoA did not affect dendritic branching and spinogenesis in scramble control neurons due to overall inactive RhoA state (*2*). As shown earlier, tomosyn knockdown neurons exhibited less dendritic complexity measured by total dendrite length (scramble: 1092.62 ± 65.77 μm, shTomosyn: 742.84 ± 50.23 μm), branch number (scramble: 23.4 ± 1.5, shTomosyn: 16.2 ± 1.0), and spine density (scramble: 0.46 ± 0.03 μm^-1^, shTomosyn: 0.33 ± 0.02 μm^-1^). When T19N-RhoA was co-expressed, tomosyn knockdown neurons showed similar total dendrite length (Fig. 5C-E) (scramble: 1163.67 ± 39.92 μm, shTomosyn: 1067.38 ± 49.17 μm), branch number (scramble: 21.5 ± 1.0, shTomosyn: 20.0 ± 1.1) and spine density (scramble: 0.47 ± 0.03 μm^-1^, shTomosyn: 0.53 ± 0.03 μm^-1^) compared to control neurons. In contrast, overexpression of WT-RhoA did not affect the compromised dendritic morphology of tomosyn knockdown neurons (Fig. 5C-E) (total dendrite length—scramble: 1169.43 ± 64.79 μm, shTomosyn: 774.95 ± 48.83 μm; branch number—scramble: 20.7 ± 1.1, shTomosyn: 15.3 ± 0.7; spine density—scramble: 0.48 ± 0.02 μm^-1^, shTomosyn: 0.34 ± 0.02 μm^-1^). Treating cultured neurons with C3T, a RhoA inhibitor, also fully rescued the dendritic phenotypes in tomosyn knockdown neurons (fig. S4A-E) (total dendrite length—scramble: 938.83 ± 41.6 μm, shTomosyn: 901.65 ± 49.91 μm, branch number—scramble: 19.7 ± 1.1, shTomosyn: 19.5 ± 1.3; spine density—scramble: 0.53 ± 0.03 μm^-1^, shTomosyn: 0.52 ± 0.02 μm^-1^). These data further suggest that tomosyn regulates dendritic arborization and the stability of dendritic spines through RhoA inhibition.

**Fig. 5.**
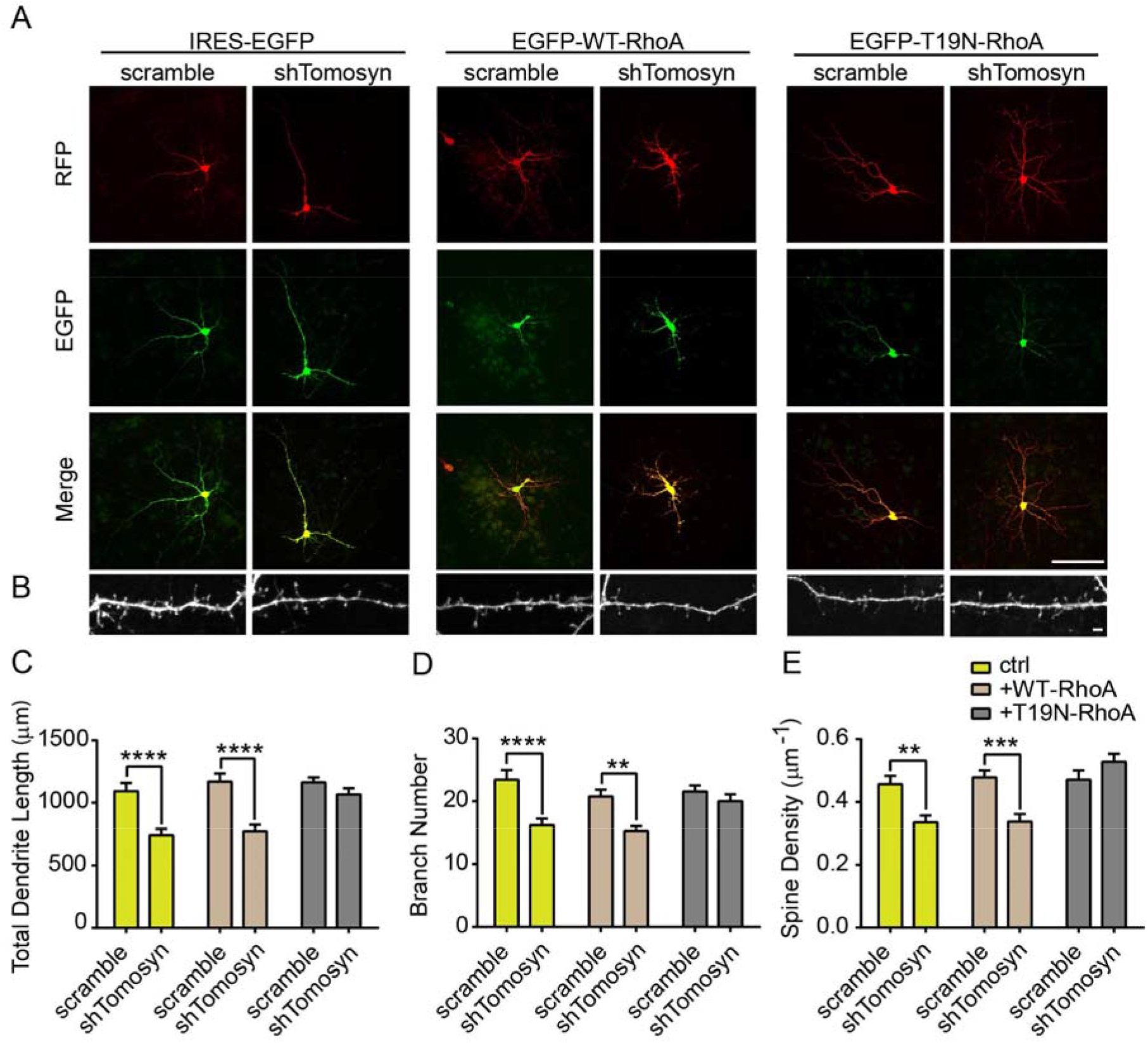
Inhibition of the Rho signaling pathway restores altered dendritic structures in tomosyn knockdown neurons. Representative images show (**A**) dendritic morphology at DIV7 and (**B**) spine morphology at DIV15. Transfected neurons were co-expressed by scramble shRNA or IRES-EGFP together with IRES-EGFP (left), EGFP-WT-RhoA (middle), and EGFP-T19N-RhoA (right). Scale bar, 100 μm in **A** and 2 μm in **B**. Quantification of (**C**) total dendrite length, (**D**) branch number, and (**E**) spine density of neurons co-transfected neurons. Co-expression of T19N-RhoA resulted in similar levels of dendritic complexity and spine density between scramble control-and shTomosyn-expressing neurons. \*\**p* < 0.01, \*\*\**p* < 0.001, \*\*\*\**p* < 0.0001 by two-way ANOVA with Sidak’s multiple comparisons test. n = 27 ∼ 35 neurons per condition in **C** and **D**; n = 18 ∼ 26 neurons per condition in **E**.

### Knockdown of tomosyn leads to altered surface-expressed glutamate receptors

Altered mEPSC kinetics in tomosyn knockdown neurons suggested that the composition of AMPA receptors may be affected (Fig. 3). We measured the surface expression of AMPA receptors by quantifying the fluorescence of pHluorin-GluR1 on the neuronal surface, including whole dendritic segments and individual spines (Fig. 6A). Knocking down tomosyn led to decreased fluorescent signals from pHluorin-GluR1 on whole dendritic surfaces (Fig. 6B), but not on individual spines (Fig. 6C). This prompted us to hypothesize that the reduced surface expression of AMPA receptors was due to the loss of dendritic spines, where most AMPA receptors are inserted. Indeed, C3T treatment significantly restored surface expression of GluR1 on dendritic segments in tomosyn knockdown neurons back to the control levels (Fig. 6B). C3T did not change the surface expression of GluR1 on spines (Fig. 6C).

**Fig. 6.**
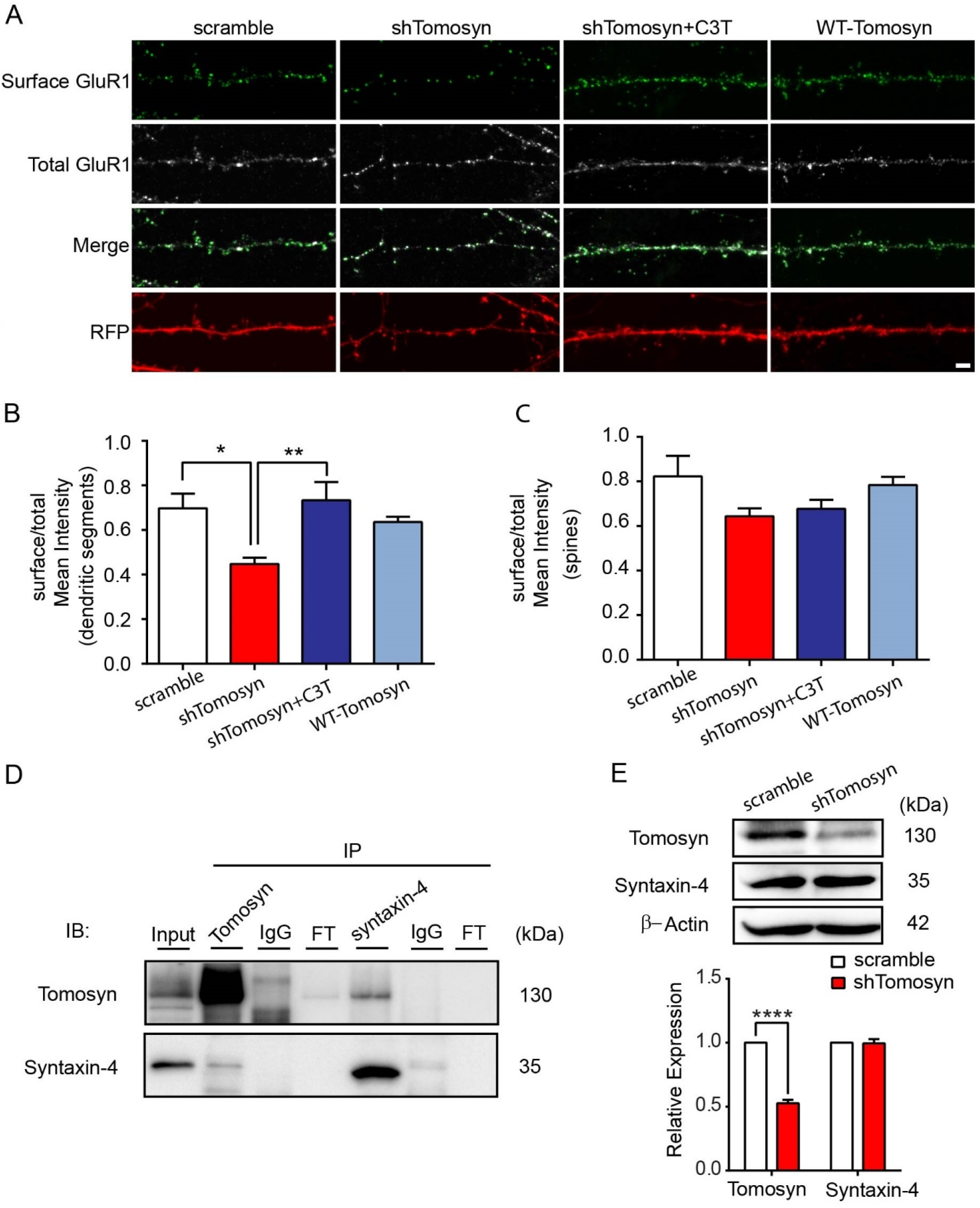
Surface-expression of GluR1 subunits is reduced in tomosyn knockdown neurons. (**A**) Representative confocal images showing the surface staining of the GluR1 subunit of AMPA receptors in cultured hippocampal neurons at 15 DIV. Neurons were co-transfected with scrambled shRNA, shTomosyn or WT-Tomosyn and pHluorin-GluR1. Tomosyn knockdown neurons were treated with the RhoA inhibitor, C3T. Scale bar, 2 μm. Surface expression of pHluorin-GluR1 were quantified via two different areas: dendritic segments (**B**) and spines (**C**). \**p* < 0.05, \*\**p* < 0.01 by one-way ANOVA with Dunnett’s multiple comparisons test. n = 20 ∼ 25 neurons per condition from three independent experiments. (**D**) Western blot analysis of co-immunoprecipitation experiments in cultured cortical neurons. Cultured cortical neuron lysates at 15 DIV were immunoprecipitated with either anti-tomosyn antibody, rabbit IgG control, anti-syntaxin-4 antibody, or mouse IgG control. Immunoblot was probed with anti-tomosyn and anti-syntaxin-4 antibodies. FT = flow through; three independent experiments. (**E**) Representative Western blot and densitometry graph show the expression of tomosyn and syntaxin-4 in N2a cells transfected with scrambled shRNA and shTomosyn. \*\*\*\**p* < 0.0001 by two-way ANOVA with Tukey’s multiple comparisons test. Data are representative of five independent cultures.

Previous studies have shown that tomosyn forms a complex with syntaxin-4 in non-neuronal cells to mediate exocytosis and inhibit mast cell degranulation (*34–36*). Syntaxin-4, as a postsynaptic t-SNARE, has been shown to regulate AMPAR trafficking in long-term potentiation (LTP) (*37*). We questioned whether tomosyn could interact with syntaxin-4 in neurons and regulate the surface expression of AMPA receptors. Co-immunoprecipitation of endogenous tomosyn and syntaxin-4 in cultured cortical neurons showed that these two proteins interact with each other in neurons (Fig. 6D). Knockdown of tomosyn, however, did not change the protein level of syntaxin-4 (Fig. 6E). Furthermore, tomosyn overexpression did not affect the surface expression of GluR1 either on dendritic segments or spines (Fig. 6A-C). These data suggested that tomosyn may regulate the surface expression of AMPA receptors through the RhoA signaling pathway rather than the SNARE machinery.

### Autism-associated tomosyn variants fail to restore dendritic complexity and the surface expression of GluR1

We have shown that the N-terminal WD40 domain of tomosyn likely inhibits RhoA activity (Fig. 4E). Two tomosyn variants harboring missense mutations (L412V and Y502C) in the WD40 domain have been identified in individuals with autism (*28*) (Fig. 7A). To evaluate potential alterations from these two proteins, shRNA-resistant tomosyn variants, L412V^r^-Tom-GFP and Y502C^r^-Tom-GFP were first engineered. When co-expressed with shTomosyn, WT^r^-Tom-GFP, L412V^r^-Tom-GFP, and Y502C^r^-Tom-GFP all effectively rescued the protein level of tomosyn (fig. S5A). RhoA activity was then determined using FRET biosensor analysis. A significant increase in FRET signals was detected in neurons expressing L412V or Y502C variant compared to those expressing WT tomosyn (fig. S5B). Neuronal morphology was further examined to determine the contributions of the shRNA-resistant tomosyn variants, L412V^r^-Tom-GFP and Y502C^r^-Tom-GFP, in tomosyn knockdown neurons. As previously shown, knockdown of tomosyn caused a reduction of dendritic arbors in total length (scramble: 1348.64 ± 104.85 μm, shTomosyn: 771.38 ± 48.58 μm) and branch number (scramble: 24.33 ± 1.6, shTomosyn: 17.09 ± 1.3) in comparison with scramble controls. WT^r^-Tom-GFP rescued the decrease in total dendrite length (1252.15 ± 60.11 μm) and branch number (22.84 ± 1.2), whereas cells co-expressing with L412V^r^-Tom-GFP or Y502C^r^-Tom-GFP still showed simplified dendritic morphology: total length (L412V: 960.7 ± 51.31 μm, Y502C: 865.03 ± 53.11 μm), branch number (L412V: 21.38 ± 1.3, Y502C: 18.53 ± 1.4) compare to scramble controls (Fig. 7B-D). Similarly, a decrease in dendritic spine density was observed as previously shown in tomosyn knockdown neurons (0.37 ± 0.02 /μm) compared to control neurons (0.61 ± 0.02 /μm) (Fig. 7E and F). Co-expression of WT^r^-Tom-GFP (0.53 ± 0.02 /μm) rescued spine loss compared to tomosyn knockdown neurons. Co-expression of Y502C^r^-Tom-GFP (0.46 ± 0.02 /μm) also showed partial spine rescue when compared to the knockdown condition. However, co-expression of L412V^r^-Tom-GFP (0.53 ± 0.02 /μm) or Y502C^r^-Tom-GFP both failed to rescue the spine loss back to the control level. Similarly, the reduced surface expression of GluR1 was restored by the WT^r^-Tom-GFP, but neither by L412V^r^-Tom-GFP nor Y502C^r^-Tom-GFP (Fig. 7G and H). Collectively, our findings support a model wherein tomosyn or ASD-associated tomosyn mutants affects dendritic arborization, spinogenesis and AMPA receptor trafficking via the RhoA signaling pathway, leading to alterations in neuronal activity (Fig. 8A-B).

**Fig. 7.**
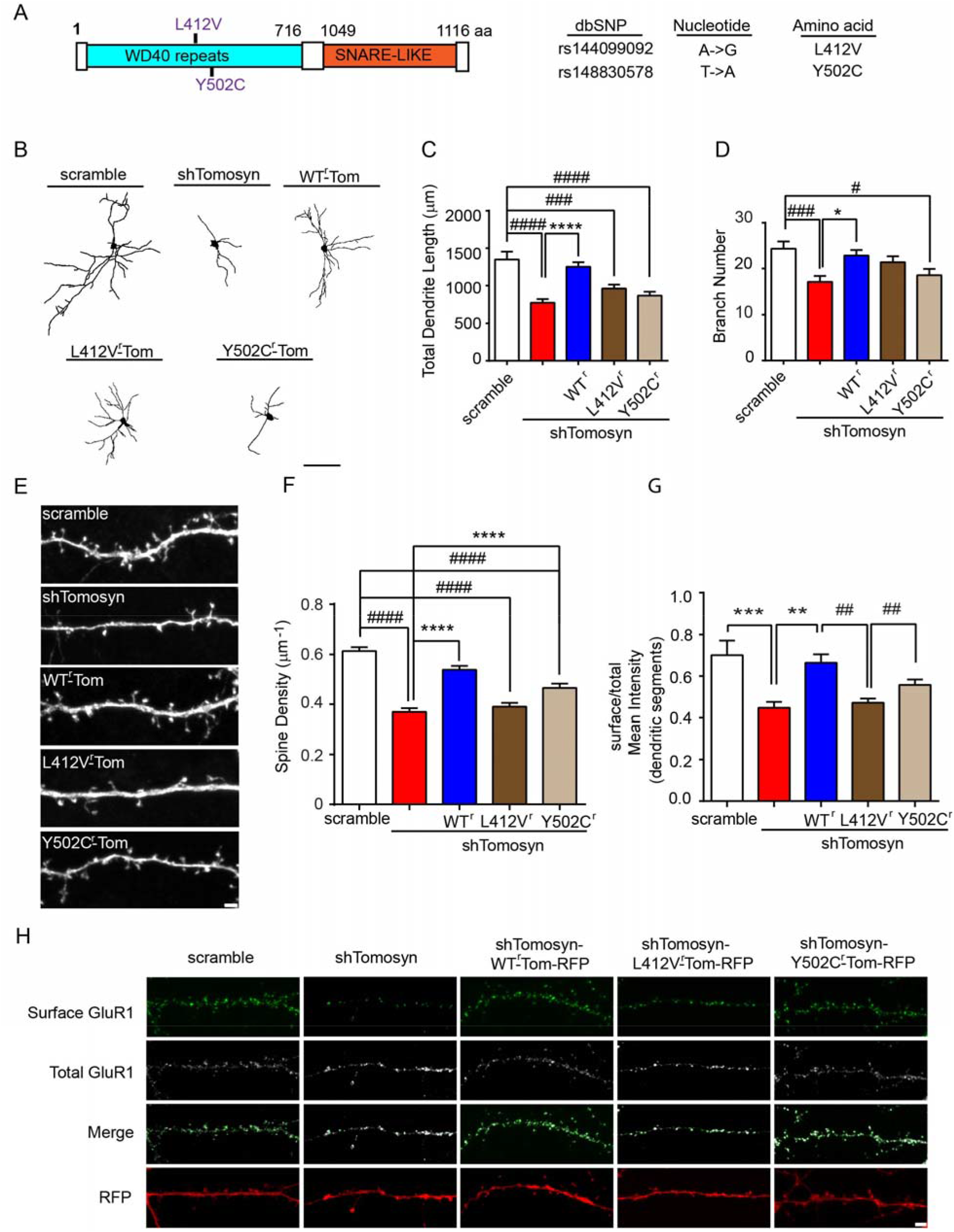
Autism-associated mutant tomosyn fails to restore dendritic arborization and spine loss in tomosyn knockdown neurons. (**A**) An illustration of the location of L412V and Y502C mutations in tomosyn. (**B**) Representative images of reconstructed dendritic trees, (**C**) total length, and (**D**) branch number from mouse hippocampal neurons transfected with scrambled shRNA + GFP, shTomosyn + GFP, shTomosyn + WT^r^-Tom-GFP, shTomosyn + L412V^r^-Tom-GFP, or shTomosyn + Y502C^r^-Tom-GFP. Scale bar, 100 μm. Both L412V and Y502C mutant tomosyn failed to rescue total dendrite length compared to scrambled shRNA. Only Y502C failed to rescue branch number compared to scrambled shRNA (^#^*p* < 0.05, ^###^*p* < 0.001, ^####^*p* < 0.0001 compared to scramble controls; \**p* < 0.05, \*\*\**p* < 0.001, \*\*\*\**p* < 0.0001 compared to shTomosyn by one-way ANOVA with Dunnett’s multiple comparisons test). n = 26 ∼ 34 neurons per condition from three independent experiments. (**E**) Representative confocal images and (**F**) the quantification of spine density in cultured hippocampal neurons transfected with different tomosyn constructs. Neurons were co-transfected with scrambled shRNA and GFP, shTomosyn and GFP, shTomosyn + WT^r^-Tom-GFP, shTomosyn + L412V^r^-Tom-GFP, or shTomosyn + Y502C^r^-Tom-GFP. Scale bar, 2 μm. ^####^*p* < 0.0001 compared to scramble controls; \*\*\*\**p* < 0.0001 compared to shTomosyn by one-way ANOVA with Dunnett’s multiple comparison tests. n = 29 ∼ 31 neurons per condition from three independent experiments. (**G**) A reduction of GluR1 surface expression in dendritic segments in neurons co-expressing shTomosyn-L412V^r^-Tom-RFP, compared to shTomosyn-WT^r^-Tom-RFP, was observed. (\*\**p* < 0.01, \*\*\**p* < 0.001 compared to shTomosyn; ^##^*p* < 0.01 compared to shTomosyn-L412V^r^-Tom-RFP by one-way ANOVA with Dunnett’s multiple comparisons test). n =17∼ 23 neurons per condition from three independent experiments. (**H**) Example confocal images show the surface expression of GluR1 in neurons expressing shTomosyn-WT^r^-Tom-RFP, shTomosyn-L412V^r^-Tom-RFP or shTomosyn-Y502C^r^-Tom-RFP together with pHluorin-GluR1. Scale bar, 2 μm.

**Fig. 8.**
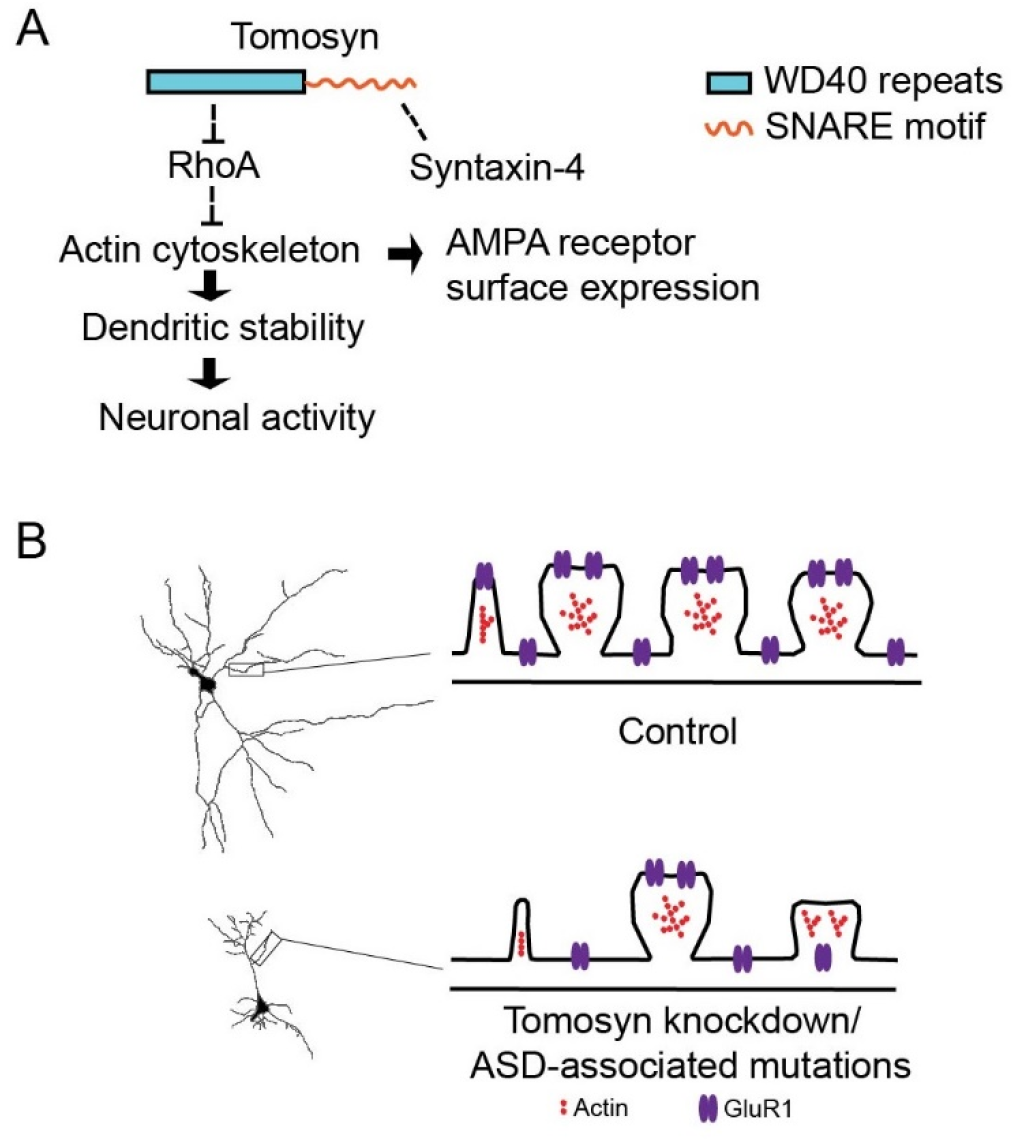
Working model of tomosyn function in regulating dendritic stability. **(A)** Mechanistic regulation of tomosyn in the maintenance of dendritic stability and AMPA receptor trafficking. Tomosyn regulates dendritic stability and AMPA receptor trafficking via inhibition of the RhoA signaling pathway. The dendritic SNARE complex formed by tomosyn and syntaxin-4 may not be required for regulating AMPA receptor trafficking. (**B**) Schematic model of spine loss, GluR1 trafficking impairment in tomosyn knockdown or autism-associated mutants rescued neurons.

## Discussion

The role of tomosyn in regulating neurotransmitter release and exocytosis has been extensively studied (*12, 35, 38, 39*). Because tomosyn is also distributed throughout dendrites (*7, 36, 40*), an undiscovered role in postsynaptic function is likely present. Here, we described the presence and function of tomosyn in postsynaptic areas. We have shown that tomosyn is a molecular link between neuronal stability and transmission, which acts as an inhibitory regulator of RhoA GTPase and plays a positive role in AMPA receptor trafficking in neurons. Autism-associated tomosyn variants L412V and Y502C showed a loss-of-function effect, resulting in a failed rescue of spine loss and simplified dendritic arborization accompanied by decreased GluR1 surface expression in tomosyn knockdown neurons.

Using biochemical approaches, we revealed that tomosyn is widely expressed in multiple brain regions, including the cerebral cortex, hippocampus, thalamus, striatum, and cerebellum, as well as diverse neuronal types, including CaMKIIα-positive excitatory neurons and GAD67-positive inhibitory neurons. Consistent with previous studies examining mRNA expression, the protein level of tomosyn is developmentally regulated and peaks at P14 in mouse brain, which is equivalent to early childhood in humans (*41, 42*). These findings suggest that tomosyn may play an important role in brain development. Our further subcellular examinations demonstrated that tomosyn is detected in both PSD fractions and in presynaptic compartments. Moreover, this study, leveraging the sparsely transfected neurons that receive unchanged presynaptic inputs, has revealed the postsynaptic role of tomosyn in neuronal morphogenesis and activity. The changes in neuronal morphology and neurotransmission upon tomosyn knockdown also showed that tomosyn exhibits a cell-autonomous effect on dendritic structures and synaptic activity. Our findings demonstrate the detailed expression profile of tomosyn in the developing brain and indicate an essential role for tomosyn in the maintenance of dendritic arborization, spine density, and synaptic strength.

Previous studies have indicated that tomosyn, containing WD40 repeats, shows substantial sequence similarity to lethal (2) giant larvae (L2gl) and yeast protein Sro7p (*43, 44*). The WD40 domain is a scaffolding structure involved in small GTPase-mediated protein-protein interactions (*45*). Interaction of Sro7p and Sec4p Rab GTPase is implicated in Rab GTPase-dependent vesicle clustering and tethering (*46, 47*). Sro7p has also been reported to disrupt cell polarity via suppression of a small GTPase Rho3p (*48*). Similar to the yeast homolog, tomosyn has recently been shown to interact with the GTP-bound state of Rab3A to partition synaptic vesicle pools (*49*). Together, this evidence suggests that tomosyn may interact with the small Rho GTPase RhoA via the WD40 domain. In the current study, FRET analysis revealed an increase of RhoA activity in tomosyn knockdown neurons. Domain analysis further indicated that the N-terminus of tomosyn with its WD40 repeats is likely responsible for mediating this RhoA inhibition. However, more sophisticated biochemical studies are required for further understanding of this inter-molecular regulation.

The dendritic SNARE fusion machinery has been implicated in the regulation of constitutive and activity-dependent glutamate receptor trafficking. Syntaxin-4 has previously been shown to regulate AMPAR trafficking in LTP, and surface expression of GluR2 and GluN1 in basal neurotransmission (*37, 50*). Synaptosomal-associated proteins (SNAPs) are members of the plasma membrane-associated SNARE family required for glutamate receptor insertion. SNAP-25, heavily studied for its role in neurotransmitter release, has recently been shown to be involved in AMPAR insertion via formation of SNAP-25-syntaxin1A/B-VAMP2 complexes (*51*). Tomosyn was first identified as a syntaxin-1-binding protein that negatively regulated neurotransmitter release and exocytosis via interaction with SNAP-25 and syntaxin-1 (*10, 52*), while later experiments in non-neuronal cells also showed that tomosyn regulates mast cell degranulation by switching interaction between syntaxin-4 and syntaxin-3 (*36*). This evidence suggests that tomosyn may play a role in glutamate receptor trafficking in hippocampal neurons via interaction with syntaxins and SNAPs. Our biochemical data showed that tomosyn indeed interacts with syntaxin-4 in neuronal cells (Fig. 6). Knocking down tomosyn might conceivably have increased the surface insertion of glutamate receptors through loss of inhibition of the dendritic SNARE complex formation. However, we found decreased GluR1 surface expression on dendritic segments in tomosyn knockdown neurons instead. Overexpression of tomosyn did not affect GluR1 surface expression neither. The reduced surface GluR1 expression in tomosyn knockdown neurons may result from decreased number of dendritic spines. A stable cytoskeletal structure is tightly modulated by de-polymerization and polymerization of actin, providing scaffolds to facilitate AMPAR trafficking along actin filaments (*53–55*). The increased RhoA activity in tomosyn knockdown neurons likely collapses the actin cytoskeleton, thereby disrupting the structure that supports AMPAR trafficking. C3T treatment, therefore, restores spine stability and consequently rescues surface expression of AMPA receptors. Our data suggest that the reduced surface expression of AMPA receptors is likely secondary to the increase of RhoA activity in tomosyn knockdown neurons, but not through loss of the interaction between tomosyn and syntaxin-4.

Although *STXBP5* is not a prominent autism-risk gene, variants have been found in individuals with ASD (*28*). In the present study, we examined two variants that carry single nucleotide mutations (L412V and Y502C) on the WD40 domain of tomosyn. ASD often features cellular characteristics involving altered dendritic structures and neuronal activity (*15, 56*). Since the WD40 domain of tomosyn likely mediates regulation of RhoA activity, L412V and Y502C variants may affect the dendritic structures of neurons by interrupting RhoA signaling. FRET analysis confirmed that neither L412V nor Y502C variant sufficiently suppressed the increased RhoA activity in tomosyn knockdown neurons. Both variants failed to completely restore total dendrite length and spine density following tomosyn knockdown and subsequently failed to restore the decreased GluR1 surface expression. All these data suggest that L412V and Y502C mutations result in the loss-of-function of tomosyn and abrogate normal RhoA signaling. It is important to note that our strategy of examining tomosyn variants in-tomosyn knockdown neurons directly reveals the functional alterations caused by the mutations relative to wild-type tomosyn, this does not mimic the likely biology of ASD. High genetic heterogeneity is a key feature for ASD, as most individuals suffer from multiple heterozygous genetic mutations. Whether L412V and Y502C variants contribute to pathological phenotypes in a more in vivo-relevant environment still requires further investigation. Nonetheless, the signaling mechanism that we described here may potentially underlie the phenotype of disrupted neuronal structures often observed in ASD brains (Fig. 8B).

Taken together, this study uncovers a crucial role of tomosyn in the maintenance of neuronal morphology and neuronal transmission via inhibition of RhoA activity. Our findings provide insights into the function of tomosyn as a regulator for maintenance of neuronal stability and activity and suggest how this mechanism may be disrupted in ASD.

## Materials and Methods

### Animals

C57BL/6J mice were obtained from the University of Maryland School of Medicine Program in Comparative Medicine. Mice were housed and cared for by the AAALAC accredited program of the University of Maryland School of Medicine. Neonatal mice of both sexes were euthanized for neuronal culture preparation. Hippocampus and whole brain from mice (2 pooled hippocampus and 1 whole brain per age group) of both sexes at different postnatal days (P) P0, P7, P14, and P21 were dissected for Western blot analysis of the temporal expression of tomosyn. Cortex, hippocampus, striatum, and cerebellum from P14 mice (2-3 pooled brain areas per sample) of both sexes were prepared for Western blot analysis of the spatial expression of tomosyn. In addition, forebrains from P21 mice (2 pooled forebrains per sample) were obtained for synaptic fraction analyses. All experiments performed were reviewed and approved by the Institutional Animal Care and Use Committees (IACUC) of the University of Maryland School of Medicine and the Hussman Institute for Autism.

### Plasmids

The pcDNA^TM^4/*myc*-His-m-tomosyn plasmid was a gift from Dr. Sushant Bhatnagar, University of Alabama. The tomosyn-GFP was engineered by adding a green fluorescent protein (GFP)-tag at the 3’ end of the pcDNA^TM^4/*myc*-His-m-tomosyn. Short hairpin (shRNA) constructs targeting tomosyn were generated by inserting the anti-tomosyn RNAi (shRNA482 also referred to as shTomosyn-GCACTGAGCGAGGAAACTAC, shRNA1083-GGAACCATATGCTGTGGTTGT) or a scrambled shRNA control (Scramble: CAGGAACGCATAGACGCATGA) into the third generation pLL3.7-RFP vector using the HpaI and XhoI restriction sites as previously described (*9*). L412V or Y502C point mutation was introduced by site-directed mutagenesis (Agilent Technologies, QuikChange II XL Site-Directed Mutagenesis Kit, Cat# 200521) using pcDNA^TM^4/*myc*-His-m-tomosyn-GFP plasmid as a template. The shRNA-resistant tomosyn (wild-type WT^r^-Tom-GFP, L412V^r^-Tom-GFP, and Y502C^r^-Tom-GFP) constructs were made by mutating three nucleotides in the shTomosyn sequences with two PCR primers: Forward 5’-AGTGGCTCTATGTGGGCACGGAACGCGGAAACATACACATTGTTA-3’ and Reverse 5’-TAACAATGTGTATGTTTCCGCGTTCCGTGCCCACATAGAGCCACT-3’. shTomosyn-WT^r^-Tom-RFP, shTomosyn-L412V^r^-Tom-RFP, and shTomosyn-Y502C^r^-Tom-RFP were generated by inserting WT^r^-Tom, L412V^r^-Tom or Y502C^r^-Tom into the shTomosyn vector. pcDNA3-EGFP-RhoA-WT (Addgene, plasmid # 12965) and pcDNA3-EGFP-RhoA-T19N (Addgene, plasmid # 12967) were a gift from Gary Bokoch (*57*). pCI-SEP-GluR1 (Addgene, plasmid #24000) was a gift from Robert Malinow (*58*). pTriEx-RhoA FLARE.sc Biosensor WT (Addgene, plasmid # 12150) was a gift from Klaus Hahn (*32*). pCAGIG (IRES-GFP) (Addgene, plasmid # 11159) was a gift from Connie Cepko (*59*). pRFP-N1 was constructed by replacing of GFP in the pEGFP-N1 (Clontech, Catalog # 6085-1) with RFP. Full-length (FL) tomosyn and domain-mutant tomosyn with BamHI and EcoRI restriction sites were subcloned into the pRFP-N1 vector. Primer sequences for FL-Tom-RFP: Forward 5’-ATATGAATTCGCCACCATGAGGAAATTCAACATCAG-3’ and Reverse 5’-ATATGGATCCATGAACTGGTACCACTTCTTATCTTTG-3’; Tom-ΔC-RFP: Forward 5’-ATATGAATTCGCCACCATGAGGAAATTCAACATCAG-3’ and Reverse 5’-ATATGGATCCATGCCCGGGATGTGTTGTG-3’; and Tom-ΔN-RFP: Forward 5’-ATATGAATTCGCCACCATGCCTGGTGGGATC-3’ and Reverse 5’-ATATGGATCCATTTTGTATTTCAGCATCATCTCATGAG-3’.

### Cell culture and transfections

Neuro-2a (N2a, a mouse neuroblastoma cell line) cells were maintained in high glucose DMEM (Invitrogen) growth media supplemented with 1% penicillin/streptomycin (Invitrogen), 2 mM L-glutamine (Invitrogen), and 10% fetal bovine serum (Sigma-Aldrich). Primary hippocampal cultures were prepared from P0 mice, and plated at a density of 3 × 10^5^ cells/well for morphometric analysis and Förster Resonance Energy Transfer (FRET) experiments, or 3 × 10^4^ cells/well for surface staining on 12-mm coverslips in 24-well plates coated with 20 μg/ml poly-(D)-lysine (Sigma-Aldrich). Neurons were maintained in serum-free neural basal media (Invitrogen) containing 1% pen/strep, 2 mM L-glutamine, and 2% B27-supplement. Lipofectamine 3000 reagent (Invitrogen, Cat# L3000075) was used to transfect N2a cells according to the manufacturer’s protocol. Primary hippocampal neurons were transfected using Lipofectamine 3000 at 4 days in vitro (DIV) followed by fixation at 7 DIV for quantifying the dendritic arborization, and calcium-phosphate transfection method at 11-12 DIV followed by fixation at 15 DIV for spine quantification. For inhibiting RhoA activity, 24 hours after transfection, neurons were either treated with 1.0 μg/mL of RhoA inhibitor C3 transferase (Cytoskeleton, Cat# CT04) for 16 hours, or 10 μL of ddH_2_O as a control for 16 hours prior to fixation. For electrophysiological recordings, primary hippocampal neurons were transfected at 9 DIV using Lipofectamine 2000 (Invitrogen, Cat# 11668-027) and incubated until 14-16 DIV.

### Synaptic fractionation

Synaptic fractionation was prepared as described previously (*60*). Briefly, forebrains were homogenized in 4 ml of 0.32 M HEPES-buffered sucrose solution (0.32 M sucrose, 4 mM HEPES, 0.25 mM PMSF and protease and phosphatase inhibitors (Thermo Fisher Scientific, Cat# 78442)) at 4 °C by using a motor-driven glass-Teflon tissue homogenizer. Homogenates (total) were centrifuged at 900 × g for 10 minutes at 4 °C to generate the nuclear fraction pellet (P1) and the supernatant (cytosol/membranes, S1). S1 was centrifuged at 10,000 × g for 15 minutes at 4 °C to obtain the crude synaptosomes (P2) and the supernatant (cytosol/light membranes, S2). P2 was washed with 0.32 M HEPES-buffered sucrose solution and re-centrifuged at 10,000 × g for 15 minutes at 4 °C. P2’, obtained from P2, was lysed in 4 ml ddH_2_O by hypoosmotic shock and was adjusted back to 4 mM HEPES. P2’ lysate was centrifuged at 25,000 × g for 20 minutes at 4 °C to isolate the synaptosomal membrane fraction (synaptosomes, P3) in the pellet and the crude vesicular fraction in the supernatant (synaptic vesicles, S3). P3 in 1 ml of 0.32 M HEPES-buffered sucrose solution was loaded on discontinuous sucrose gradient containing 3.0 ml of 0.8 M sucrose, 3.0 ml of 1 M sucrose and 3.5 ml of 1.2 M sucrose in 4 mM HEPES-buffered solution, and re-centrifuged at 150,000 × g for 2 hours at 4 °C to obtain synaptic plasma membranes (SPM). SPM was re-suspended in Triton X-100/HEPES/EDTA solution (0.54% Triton X-100, 50 mM HEPES, 2 mM EDTA) and centrifuged at 32,000 × g for 20 minutes at 4 °C to obtain the post-synaptic density fraction (PSD). A quantity of 10 μg protein from each fractionation was loaded for Western blot analysis.

### Western blot analysis

Brain tissues or N2a cells transfected with plasmids were harvested in RIPA buffer containing 20 mM Tris-HCl (pH 7.5), 150 mM NaCl, 1 mM Na_2_EDTA, 1 mM EGTA, 1% NP-40, 1% sodium deoxycholate, 2.5 mM sodium pyrophosphate, 1 mM beta-glycerophosphate, 1 mM Na_3_VO_4_, 1 μg/ml leupeptin, 1 mM PMSF, and a phosphatase and protease inhibitor cocktail (Sigma-Aldrich, Cat# MSSAFE). Protein concentration was determined using a Pierce BCA Protein Assay Kit (Thermo Fisher Scientific, Cat# 23227). 50 μg of total protein from N2a cell lysates was used to determine the knockdown efficiency of tomosyn shRNAs. 10 μg of total protein from different brain regions was used to examine tomosyn expression. Protein samples were run on 8% gels for SDS-PAGE and transferred to a PVDF membrane. After blocking the membrane with 5% skim milk in TBST (0.1% Tween 20 in TBS) for 1 hour at room temperature, the membrane was immunoblotted with the following primary antibodies: rabbit anti-tomosyn (1:1000, Synaptic Systems, Cat# 183103), mouse anti-β-Actin (1:5000, Sigma-Aldrich, Cat# A1978), mouse anti-syntaxin-4 (1:1000, Synaptic Systems, Cat# 110041), mouse anti-PSD-95 (1:1000, UC Davis/NIH NeuroMab Facility, Cat# 75-028). The membrane was washed 3 times with TBST for 10 minutes and incubated with the HRP-conjugated secondary antibodies for 60 minutes at room temperature. The membrane was then washed 5 times with TBST for 10 minutes at room temperature. The chemiluminescent blots were imaged with the ChemiDoc^TM^ Touch Imaging System (Bio-Rad). The densitometry of the protein signals was analyzed using Image Lab^TM^ software (Bio-Rad).

### Immunocytochemistry and morphometric analysis

Cultured neurons were fixed at 7 or 15 DIV with 4% paraformaldehyde in phosphate buffer (PB) (320 mM Na2HPO4, 76 mM NaH2PO4) for 15 minutes and permeabilized and blocked in TBS containing 0.1% Triton X-100, 3% BSA and 1% donkey serum. Cells were immunostained with the following primary antibodies: mouse anti-GFP (1:5000, Molecular Probes, Cat# A-11120), rat anti-RFP (1:1000, Chromotek, Cat# 5f8-100), rabbit anti-tomosyn (1:500, Synaptic Systems, Cat# 183103), mouse anti-MAP2 (1:2000, Sigma-Aldrich, Cat# M9942), mouse anti-CaMKIIα (1:500, Thermo Fisher Scientific, Cat# MA-048), and mouse anti-GAD67 (1:500, Millipore Sigma, Cat# MAB5406) followed by Alexa Fluor 488, 568, 594-conjugated secondary antibodies (Invitrogen). For dendrite analysis, z-stack images were taken using a confocal laser scanning microscope (Zeiss LSM 780) with a 40 X/1.3 oil DIC objective. Original images of neurons were used to trace dendrites for dendrite length measurement using ImageJ (NIH) with NeuronJ plugin (*61*). Traced neurons were further analyzed using ImageJ with Sholl analysis plugin (*62*). Concentric circles having 10 μm increments in radius were defined from the center of the cell body. The number of traced dendrites crossing each circle was counted. For spine analysis, 2–3 dendrite segments per neuron were selected randomly from secondary branches on the apical dendrite. Z-stack images were taken using Zeiss LSM 780 with a 63 ×/1.4 oil DIC objective and 4 zooms at 512 × 512 resolution. The number of dendritic spines, which is ≤; 2 μm in length, were manually counted using ImageJ. Images of neurons co-immunostained for either CaMKIIα or GAD67 and tomosyn were taken using the EVOS FL Auto2 imaging system with a 40 ×/0.95 NA air objective (Invitrogen).

### Electrophysiology

Neurons at 14-16 DIV were used for whole-cell electrophysiological recordings following either scrambled shRNA or shTomosyn transfection at 9 DIV. Coverslips were transferred from a 24-well plate into a recording chamber superfused with a buffer containing the following (in mM): 125 NaCl, 3 KCl, 10 HEPES, 5 dextrose, 1 MgCl_2_, and 2 CaCl_2_. The recording buffer was adjusted to ∼260 mOsm and pH 7.2, and the bath was maintained at a temperature of 30°C. Glass pipettes pulled to a resistance of 3-7 MΩ were filled with internal solution containing the following (in mM): 134 KMeSO_4_, 3 KCl, 10 HEPES, 1 MgCl_2_, 4 MgATP, 0.5 Na_2_GTP, 5 K_2_-creatine phosphate, and 5 Na_2_-creatine phosphate. The internal solution was adjusted to ∼250 mOsm and pH 7.4. Only RFP-expressing cells with a series resistance ≤ 30 MΩ and a resting membrane potential ≤ −50 mV were accepted for final analysis. Whole-cell parameters were monitored throughout the recording with a 100 ms, −10 mV step delivered every 30 seconds. Recordings were made using an Axon MultiClamp 700B amplifier (Molecular Devices). Data were filtered at 2 kHz and digitized at 10 kHz with a National Instruments digital-analog converter under the control of Igor Pro software (WaveMetrics). Miniature EPSCs were recorded in voltage mode (Vh = −70 mV) in the presence of 50 μM AP5, 1 μM tetrodotoxin (TTX), and 10 μM gabazine (GBZ). The frequency, amplitude, and kinetics of mEPSCs were analyzed with Mini Analysis software (Synaptosoft), with a threshold of 3 × RMS noise for event discrimination. At least 50 events per cell were used in these calculations.

### FRET analysis

Hippocampal neurons were co-transfected with pTriEx-RhoA FLARE.sc Biosensor WT and either scrambled shRNA or shTomosyn at 4 DIV. Cells were fixed with 4% paraformaldehyde in PB and mounted at 7 DIV. Imaging was performed with an upright Zeiss confocal LSM 780 and a 40× EC Plan-NeoFluar/1.3 oil-immersion objective lens. For sensitized emission, FRET excitation was performed using the 458 nm laser, and emission filter bands were set to 440-510 for CFP and 517-606 for YFP. The 561 nm laser was used for excitation of RFP to identify shRNA-transfected neurons. Single plane images were exported to ImageJ for analysis. The RFP channel image was used as a reference to create a whole-cell ROI (region-of-interest), which was subsequently overlaid on CFP and YFP channel images and mean fluorescence intensity (I) was measured within the ROI. As the RhoA-FLARE is an intramolecular FRET biosensor, the FRET efficiency (E_FRET_) is calculated by simple ratiometry of the acceptor (YFP) over donor (CFP) intensity: E_FRET_ = IYFP/ICFP. Acceptor photobleaching FRET was done by placing a circular bleach ROI (r = 2.04 μm) over secondary or tertiary parts of either apical or basal dendritic compartments of transfected neurons. Acceptor bleaching was executed over 50 iterations with the 514 nm laser at maximum power. Changes in fluorescence intensities in the donor (CFP) and acceptor (YFP) channels were recorded using low power (2%) excitation with the 458 nm and 514 nm lasers, respectively, and a total of 10 scans were acquired: 5 before (pre) and 5 after (post) the single bleaching event. The mean fluorescence intensity values (I) of ROIs were exported to Excel. The donor channel bleach ROI data were background-corrected and normalized to unbleached background ROIs and reference ROIs, respectively. FRET efficiency (E_FRET_) in percentage was calculated according to: E_FRET_ = ((Idonor_post_ − Idonor_pre_) /Idonor_post_) × 100. All calculated E_FRET_ data was exported to GraphPad Prism 6 for statistical analysis.

### Immunoprecipitation

Immunoprecipitations of endogenous binding partners of tomosyn or syntaxin-4 were performed using lysates of cultured cortical neurons. Cultured cortical neurons were lysed in lysis buffer (20 mM Tris, pH 7.5, 150 mM NaCl, 1% Triton X-100) containing phosphatase and protease inhibitor cocktail (Thermo Fisher Scientific, Cat# 78442). Lysates were precleared with protein A/G agarose beads (Thermo Fisher Scientific, Cat# 20421) for 1 hour. 2 μg of rabbit anti-tomosyn or mouse anti-syntaxin-4 antibody was added to the lysate followed by rotation overnight at 4 °C. 20 μL of protein A/G agarose was added to the immune complex and incubated for 1 hour at 4 °C. The beads were washed five times with TBST. Bound proteins were eluted in 2 × Laemmli sample buffer (Bio-Rad, Cat# 161-0737) containing 5% 2-mercaptoethanol by heating at 95 °C for 5 minutes.

### Surface staining

Neurons were co-transfected pCI-SEP-pHluorin-GluR1 with scrambled shRNA or shTomosyn. The surface staining of pHluorin-GluR1 was performed as previously described (*51, 63*). Briefly, transfected neurons were incubated with mouse anti-GFP antibody in pre-chilled artificial cerebrospinal fluid (ACSF) (124 mM NaCl, 5 mM KCl, 1.23 mM NaH_2_PO_4_, 26 mM NaHCO_3_, 1 mM MgCl_2_, 2 mM CaCl_2_,) containing 10% BSA for 1 hour at room temperature. Cells were washed three times with ACSF, incubated with donkey anti-mouse Alex Fluor 647-conjugated secondary antibody (1:1000, Invitrogen) for 1 hour at room temperature, and again washed three times with ACSF. For total pHluorin-GluR1 staining, cells were fixed with 4% paraformaldehyde in PB for 15 minutes, permeabilized, and blocked in blocking solution. Cells were immunostained with rabbit anti-GFP (1: 4000, Antibodies-Online, Cat# ABIN346939) and rat anti-RFP antibodies. Cells were washed three times followed by incubation with goat anti-rat Alexa Fluor 568-and donkey anti-rabbit Alexa Fluor 488-conjugated secondary antibody. Coverslips were mounted on the glass slides with Prolong Diamond Antifade Mountant (Molecular Probes, Cat# P36961) prior to imaging. 2–3 dendrite segments per neuron were selected randomly from secondary branches on the apical dendrite. Identical acquisition settings were applied to each image within all experiments. For surface GluR1 expression quantification, both whole dendritic segments and individual spines are chosen as ROI. Surface GluR1/total GluR1 was calculated as surface/total = (mean intensity of surface − background intensity) / (mean intensity of total − background intensity).

### Statistical analysis

Sample sizes for each experiment are based on previously published studies from our laboratory and standards in the field. Values were expressed as mean ± SEM (standard error of the mean). n = number of neurons or repeats. All statistical comparisons were performed using GraphPad Prism 6 (GraphPad Software, San Diego, CA). Variable comparisons between groups were performed using either one-way ANOVA with Dunnett’s *post hoc* test or two-way ANOVA with Tukey’s or Sidak’s *post hoc* test. P-values smaller than 0.05 were considered significant.

## Acknowledgments

We want to thank Drs. Louis DeTolla and Turhan Coksaygan at the University of Maryland School of Medicine for providing veterinary service and consultation. The authors gratefully acknowledge Elizabeth Benevides and Drs. Gene J. Blatt and John P. Hussman for their edits and helpful comments on the manuscript. This work is supported by Hussman Foundation grant HIAS15003 to Y-C. L, and HIAS18001 to SH.

## Supplementary Materials

**Fig. S1.**
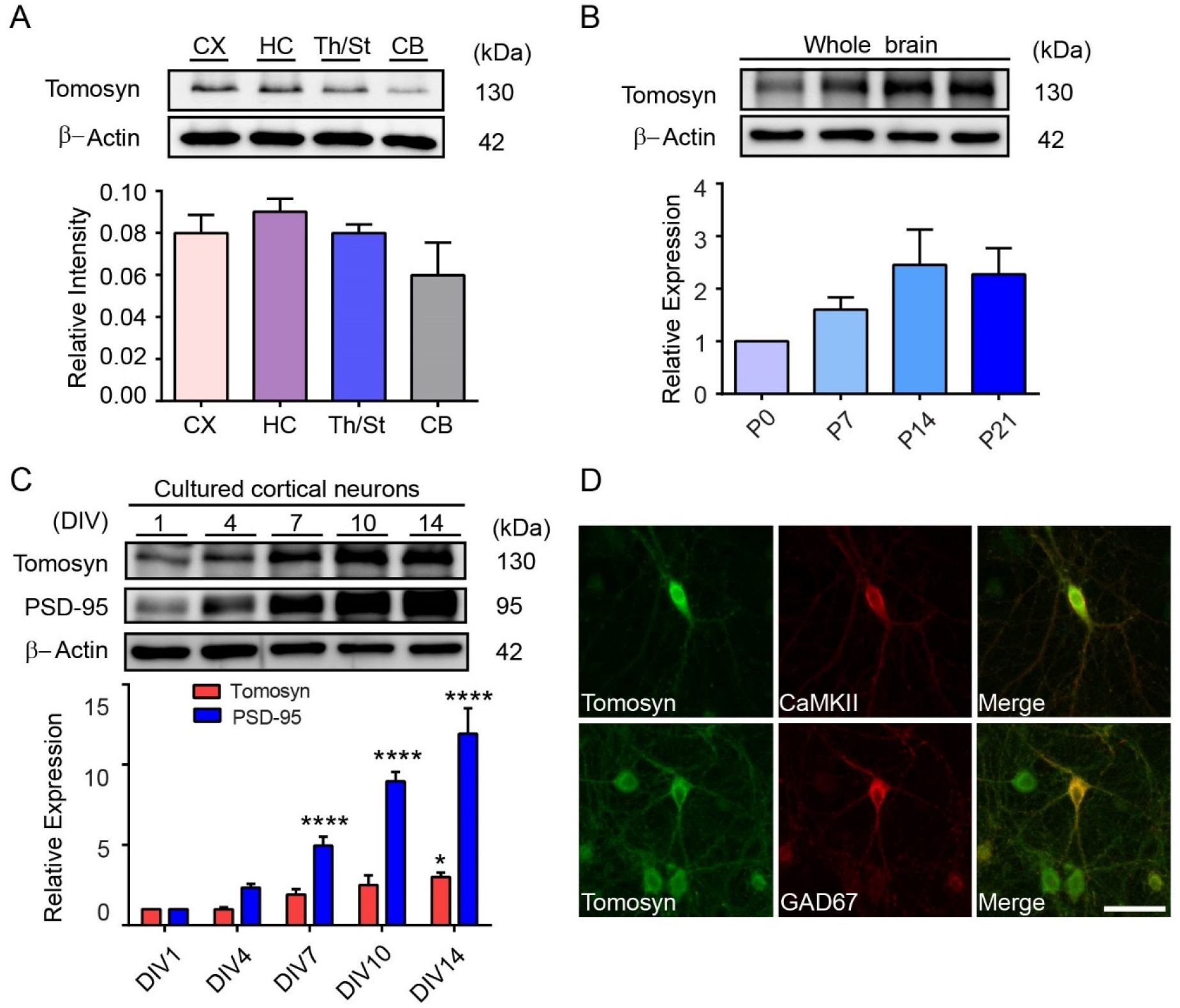
Spatial and temporal expression patterns of tomosyn during different developmental stages. (**A**) Spatial expression of tomosyn in different brain regions at P14. CX, cortex; HC, hippocampus; Th/St, thalamus and striatum; CB, cerebellum. (**B**) Temporal expression pattern of tomosyn in whole brain at P0, P7, P14, and P21. (**C**) Temporal expression of tomosyn and PSD-95 in cultured cortical neurons at 1, 4, 7, 10, and 14 DIV. \**p* < 0.05, \*\*\*\**p* < 0.0001 compared to 1 DIV by one-way ANOVA with Dunnett’s multiple comparisons test. (**D**) Fluorescent images showing either CaMKIIα or GAD67 and tomosyn from cultured hippocampal neurons at 15 DIV. Scale bar, 20 μm. Data are representative of three independent experiments in **A**-**D**.

**Fig. S2.**
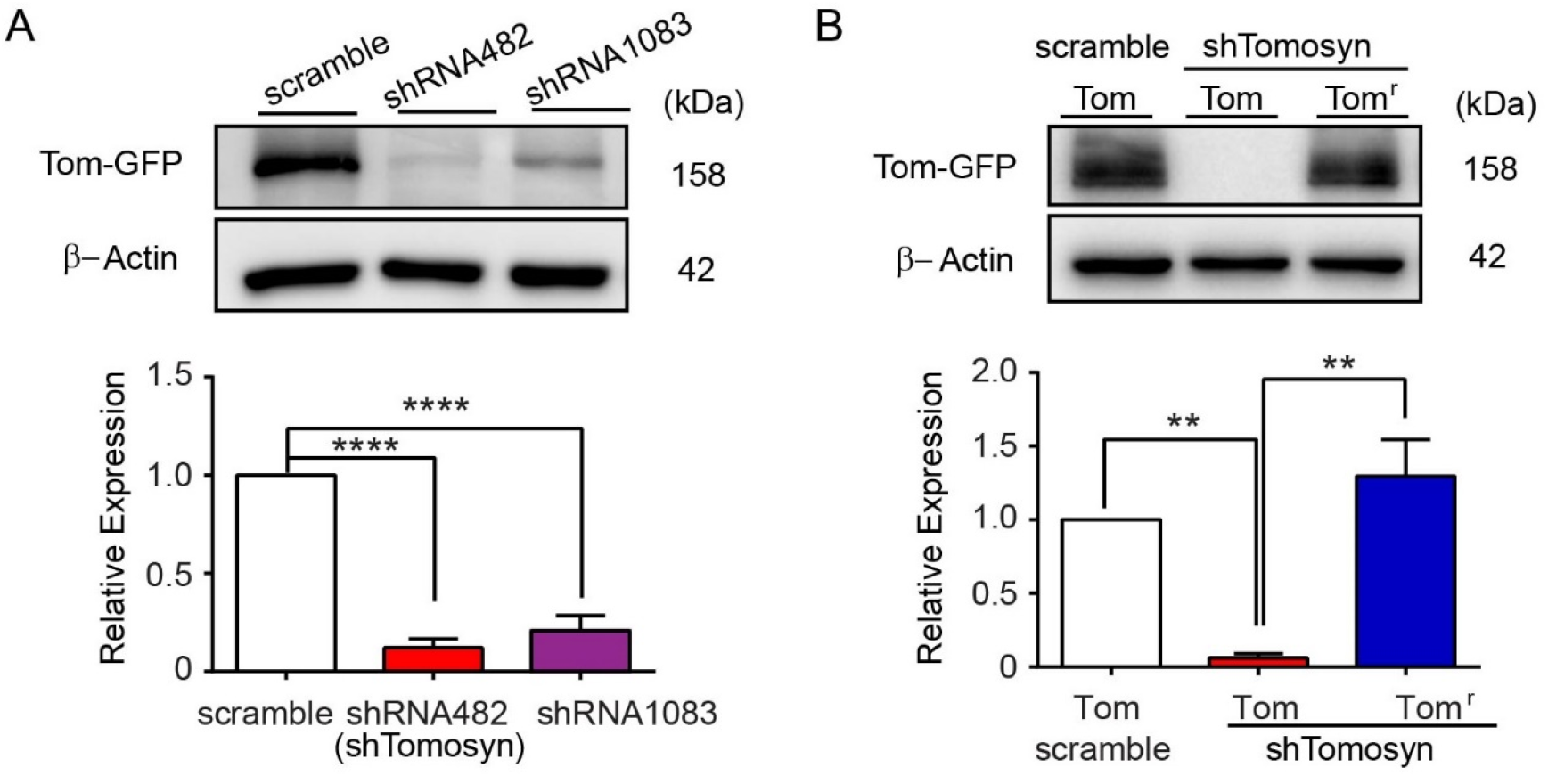
Silencing effect of shRNA on tomosyn and rescuing effect of shRNA-resistant tomosyn. (**A**) Western blot analysis shows the knockdown efficiency of shRNAs against tomosyn. Tomosyn-GFP was co-transfected with scrambled shRNA, shRNA482, or shRNA1083 in N2a cells. shRNA482 resulted in the most effective reduction of tomosyn protein level and was shown as shTomosyn in subsequent experiments. \*\*\*\**p* < 0.0001 compared to scramble control by one-way ANOVA with Dunnett’s multiple comparisons test. Data are representative of three independent experiments. (**B**) Western blot analysis shows the protein level of Tom^r^-GFP, an shRNA-resistant form of tomosyn, in tomosyn-knockdown N2a cells. While shTomosyn reduced the expression of Tom-GFP, the protein level of Tom^r^-GFP was not affected by shTomosyn knockdown. \*\**p* < 0.01 when compared with shTomosyn + Tom-GFP by one-way ANOVA with Dunnett’s multiple comparisons test. Data are representative of three independent experiments.

**Fig. S3.**
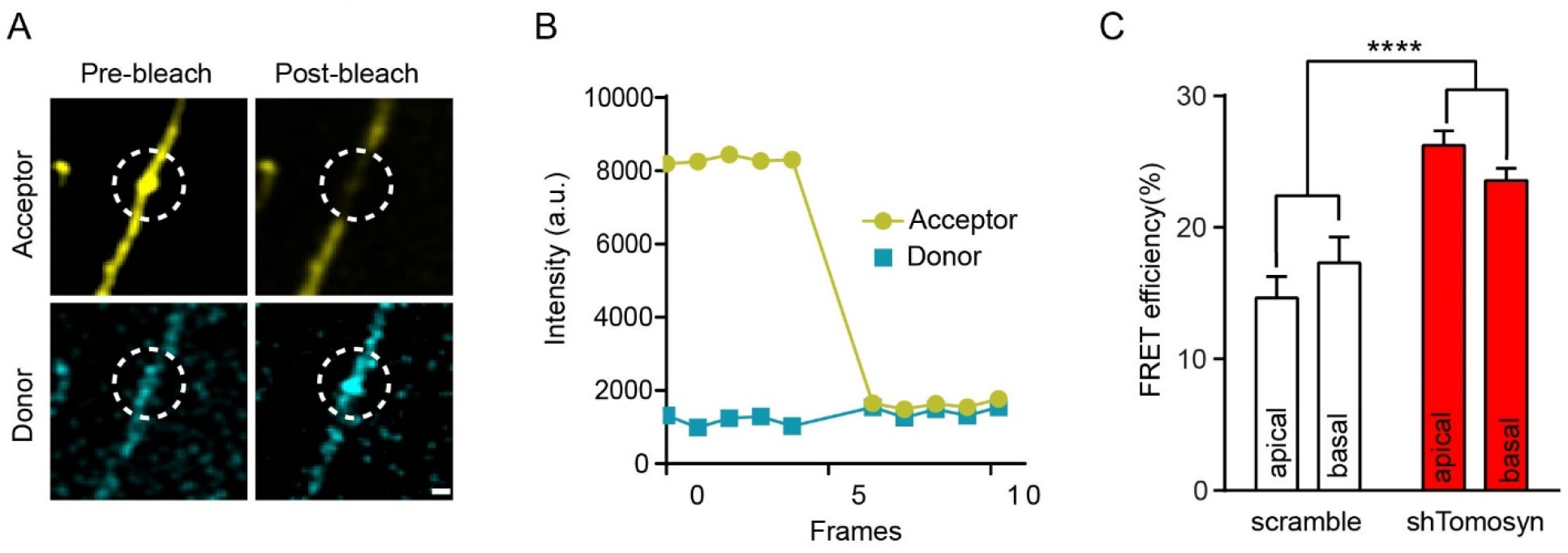
Tomosyn knockdown leads to increased RhoA activity in different dendritic areas. (**A**) Representative images of dendritic segments taken immediately before and after photobleaching. pbFRET was used to measure RhoA activity in different dendritic areas. Bleached area was highlighted as a white dashed circle (ROI). Scale bar, 2 μm. (**B**) A representative graph of a pbFRET experiment shows changes in fluorescence intensities within bleach ROI over the course of 10 frames. Photobleaching was performed after the 5th frame. (**C**) FRET efficiency was determined by calculating percentage (%) of increase in donor fluorescence intensity after acceptor photobleaching. Hippocampal neurons were co-transfected with a RhoA Biosensor and either scrambled shRNA or shTomosyn. Tomosyn knockdown neuron showed higher FRET ratio. \*\*\*\**p* < 0.0001 when compared to scramble control. No significant FRET efficiency percentage (%) of increase was found between apical dendrites and basal dendrites within each group. *p* = 0.92. n = 17 neurons per condition from four independent experiments. Factor: dendrite region. Data are analyzed by two-way ANOVA with Tukey’s multiple comparisons test.

**Fig. S4.**
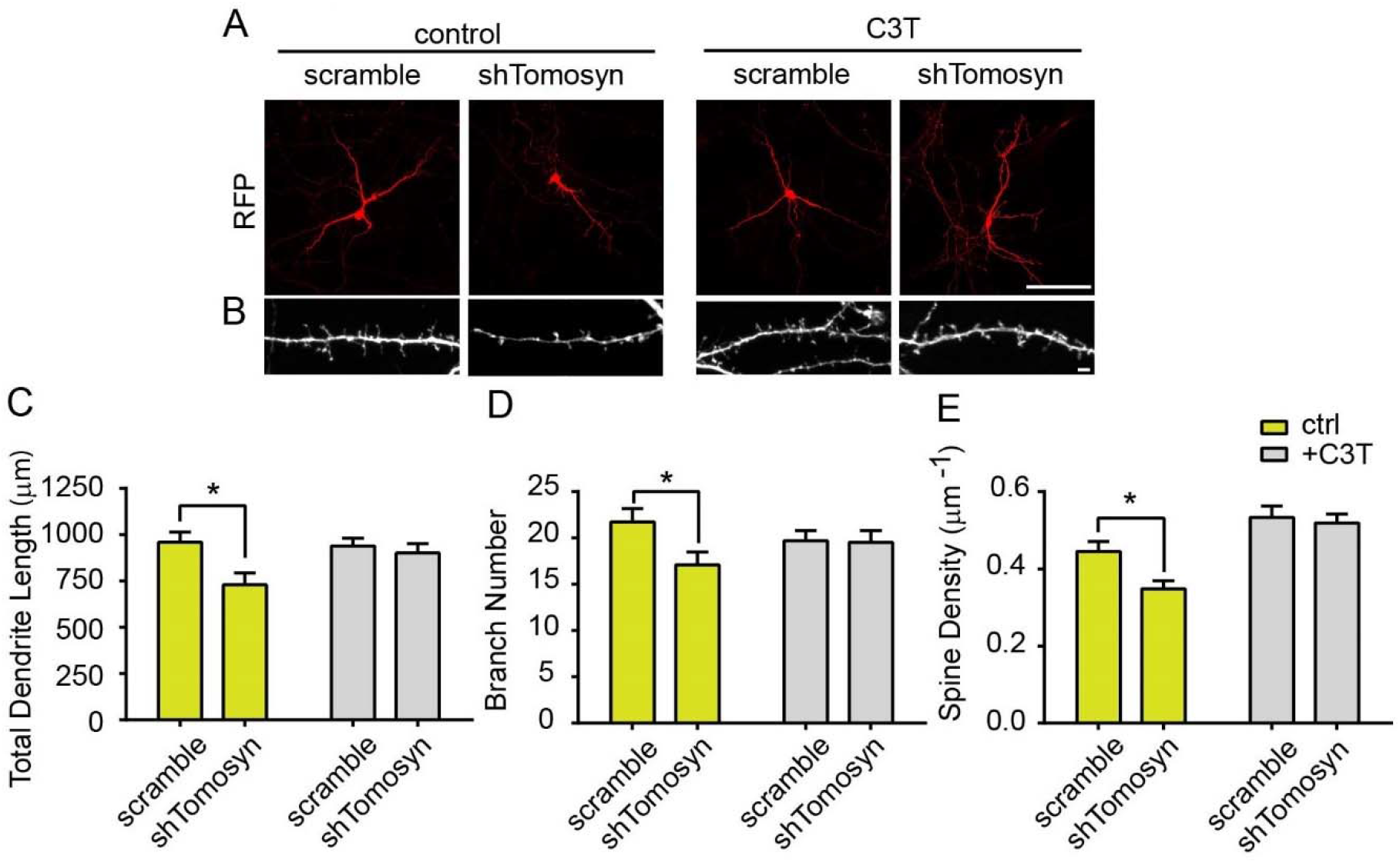
Inhibition of the Rho signaling pathway restores altered dendritic structures in tomosyn knockdown neurons. Transfected neurons expressing scramble shRNA or shTomosyn were treated with control (ddH_2_O) and C3T. (**A**) Representative images show dendritic morphology at 7 DIV. Scale bar, 100 μm. (**B**) Representative images show spine morphology at 15 DIV. Scale bar, 2 μm. Tomosyn knockdown neurons treated with C3T rescued the (**C**) total dendrite length, (**D**) branch number, and (**E**) spine density. \**p* < 0.05 by two-way ANOVA with Sidak’s multiple comparisons test. n = 23∼32 neurons per condition in **C** and **D**; n = 23 ∼ 30 neurons per condition in **E**. Data are representative of three independent experiments in **A**-**E**.

**Fig. S5.**
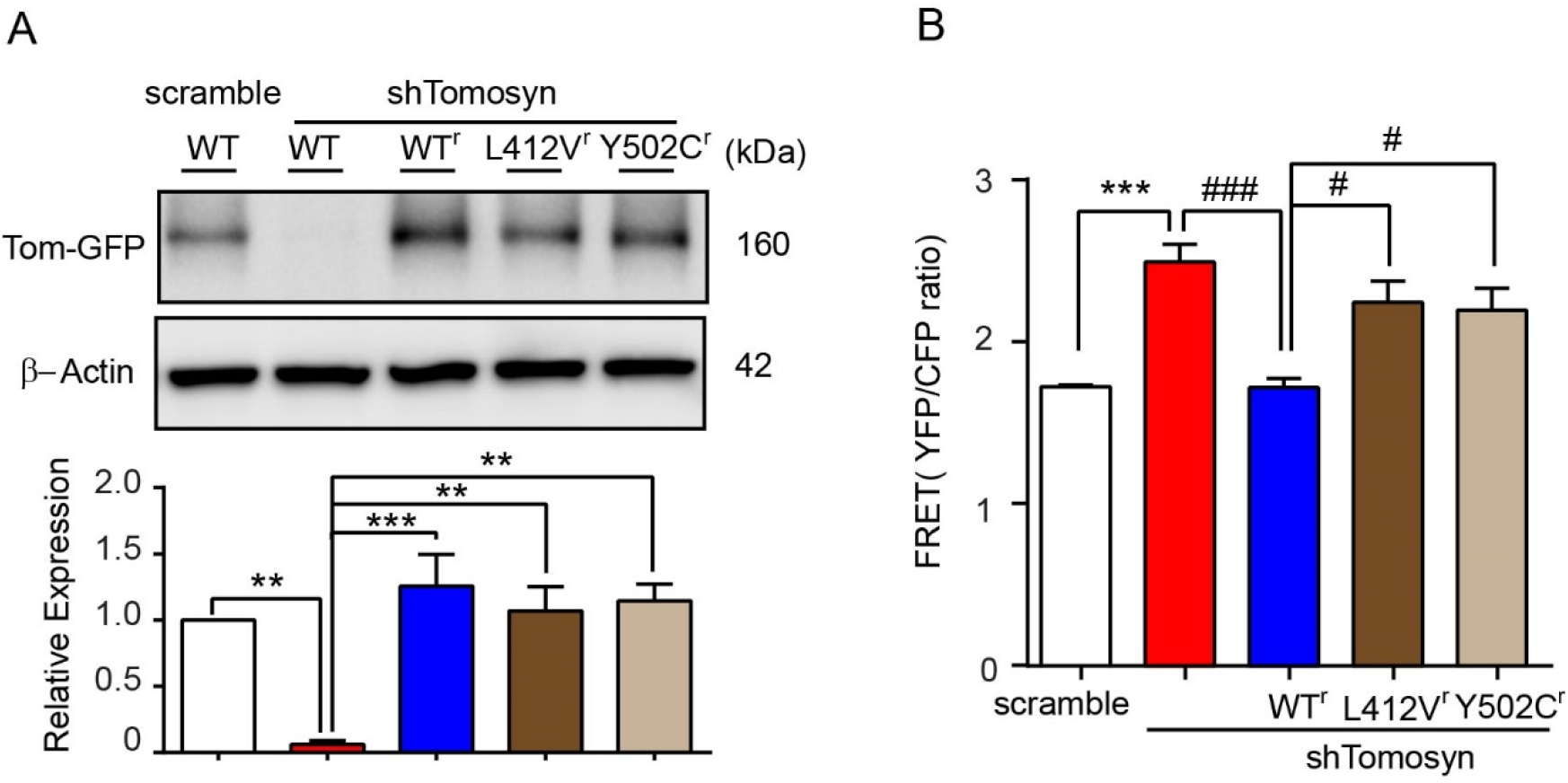
shRNA-resistant autism-associated tomosyn variants result in increased RhoA activity in tomosyn knockdown neurons. (**A**) Representative Western blot and densitometry graph show protein levels of shRNA-resistant WT^r^-Tom-GFP, L412V^r^-Tom-GFP, and Y502C^r^-Tom-GFP in N2a cells co-transfected with shTomosyn. \*\**p* < 0.01, \*\*\**p* < 0.001 by one-way ANOVA with Dunnett’s multiple comparisons test. Data are representative of three independent experiments. (**B**) Representative graph showing FRET efficiency in tomosyn knockdown neurons rescued with WT^r^, L412V^r^, or Y502C^r^ tomosyn constructs. Neurons were co-transfected with a RhoA biosensor and scrambled shRNA, shTomosyn, shTomosyn-WT^r^-Tom-RFP, shTomosyn-L412V^r^-Tom-RFP, or shTomosyn-Y502C^r^-Tom-RFP. \*\*\**p* < 0.001 compared to scrambled shRNA; ^#^*p* < 0.05, ^###^*p* < 0.001 compared to shTomosyn-WT^r^-Tom-RFP by one-way ANOVA with Dunnett’s multiple comparisons test. Data are representative of four independent experiments.

## References

1. Y. C. Lin, A. J. Koleske, Mechanisms of synapse and dendrite maintenance and their disruption in psychiatric and neurodegenerative disorders. Annu Rev Neurosci 33, 349–378 (2010).

2. A. Y. Nakayama, M. B. Harms, L. Luo, Small GTPases Rac and Rho in the Maintenance of Dendritic Spines and Branches in Hippocampal Pyramidal Neurons. J Neurosci 20, 5329–5338 (2000).

3. E. E. Govek et al., The X-linked mental retardation protein oligophrenin-1 is required for dendritic spine morphogenesis. Nat Neurosci 7, 364–372 (2004).

4. A. Reiner, J. Levitz, Glutamatergic Signaling in the Central Nervous System: Ionotropic and Metabotropic Receptors in Concert. Neuron 98, 1080–1098 (2018).

5. G. H. Diering, R. L. Huganir, The AMPA Receptor Code of Synaptic Plasticity. Neuron 100, 314–329 (2018).

6. C. G. Lau, R. S. Zukin, NMDA receptor trafficking in synaptic plasticity and neuropsychiatric disorders. Nat Rev Neurosci 8, 413 (2007).

7. Y. Fujita et al., Tomosyn: a syntaxin-1-binding protein that forms a novel complex in the neurotransmitter release process. Neuron 20, 905–915 (1998).

8. E. S. Masuda, B. C. Huang, J. M. Fisher, Y. Luo, R. H. Scheller, Tomosyn binds t-SNARE proteins via a VAMP-like coiled coil. Neuron 21, 479–480 (1998).

9. Y. Ben-Simon et al., A Combined Optogenetic-Knockdown Strategy Reveals a Major Role of Tomosyn in Mossy Fiber Synaptic Plasticity. Cell Rep 10.1016/j.celrep.2015.06.037 (2015).

10. T. Sakisaka et al., Dual inhibition of SNARE complex formation by tomosyn ensures controlled neurotransmitter release. J Cell Biol 183, 323–337 (2008).

11. B. Barak et al., Neuron-specific expression of tomosyn1 in the mouse hippocampal dentate gyrus impairs spatial learning and memory. Neuromolecular Med 15, 351–363 (2013).

12. S. R. Batten et al., Linking kindling to increased glutamate release in the dentate gyrus of the hippocampus through the STXBP5/tomosyn-1 gene. Brain Behav 7, e00795–e00795 (2017).

13. T. Sakisaka et al., Regulation of SNAREs by tomosyn and ROCK: implication in extension and retraction of neurites. J Cell Biol 166, 17–25 (2004).

14. J. J. Saldate, J. Shiau, V. A. Cazares, E. L. Stuenkel, The ubiquitin-proteasome system functionally links neuronal Tomosyn-1 to dendritic morphology. J Biol Chem 293, 2232–2246 (2018).

15. M. P. Forrest, E. Parnell, P. Penzes, Dendritic structural plasticity and neuropsychiatric disease. Nat Rev Neurosci 19, 215 (2018).

16. D. R. Hampson, G. J. Blatt, Autism spectrum disorders and neuropathology of the cerebellum. Front Neurosci 9, 420–420 (2015).

17. K. Subramanian et al., Basal ganglia and autism – a translational perspective. Autism Res 10, 1751–1775 (2017).

18. American Psychiatric Association, Diagnostic and Statistical Manual of Mental Disorders, 5th Edn. A. P. Association, Ed. (Arlington, VA: American Psychiatric Publishing: American Psychiatric Press, Inc, 2013).

19. S. J. Sanders et al., Insights into Autism Spectrum Disorder Genomic Architecture and Biology from 71 Risk Loci. Neuron 87, 1215–1233 (2015).

20. P. J. Short et al., De novo mutations in regulatory elements in neurodevelopmental disorders. Nature 555, 611–616 (2018).

21. I. Iossifov et al., The contribution of de novo coding mutations to autism spectrum disorder. Nature 515, 216–221 (2014).

22. M. Chahrour et al., Current Perspectives in Autism Spectrum Disorder: From Genes to Therapy. J Neurosci 36, 11402–11410 (2016).

23. L. Volk, S. L. Chiu, K. Sharma, R. L. Huganir, Glutamate synapses in human cognitive disorders. Annu Rev Neurosci 38, 127–149 (2015).

24. Y. C. Lin, J. A. Frei, M. B. Kilander, W. Shen, G. J. Blatt, A Subset of Autism-Associated Genes Regulate the Structural Stability of Neurons. Front Cell Neurosci 10, 263 (2016).

25. M. S. Bridi, S. M. Park, S. Huang, Developmental Disruption of GABA(A)R-Meditated Inhibition in Cntnap2 KO Mice. eNeuro 4, ENEURO.0162-0117.2017 (2017).

26. A. L. Oblak, T. T. Gibbs, G. J. Blatt, Reduced GABAA receptors and benzodiazepine binding sites in the posterior cingulate cortex and fusiform gyrus in autism. Brain Res 1380, 218–228 (2011).

27. A. L. Oblak, T. T. Gibbs, G. J. Blatt, Decreased GABA(B) receptors in the cingulate cortex and fusiform gyrus in autism. J Neurochem 114, 1414–1423 (2010).

28. H. N. Cukier et al., Exome sequencing of extended families with autism reveals genes shared across neurodevelopmental and neuropsychiatric disorders. Mol Autism 5, 1 (2014).

29. J. P. Hussman et al., A noise-reduction GWAS analysis implicates altered regulation of neurite outgrowth and guidance in autism. Mol Autism 2, 1 (2011).

30. L. K. Davis et al., Novel copy number variants in children with autism and additional developmental anomalies. J Neurodev Disord 1, 292–301 (2009).

31. N. Matsunami et al., Identification of rare recurrent copy number variants in high-risk autism families and their prevalence in a large ASD population. PloS one 8, e52239–e52239 (2013).

32. O. Pertz, L. Hodgson, R. L. Klemke, K. M. Hahn, Spatiotemporal dynamics of RhoA activity in migrating cells. Nature 440, 1069 (2006).

33. K. Aoki, M. Matsuda, Visualization of small GTPase activity with fluorescence resonance energy transfer-based biosensors. Nat Protoc 4, 1623–1631 (2009).

34. C. H. Widberg, N. J. Bryant, M. Girotti, S. Rea, D. E. James, Tomosyn interacts with the t-SNAREs syntaxin4 and SNAP23 and plays a role in insulin-stimulated GLUT4 translocation. J Biol Chem 278, 35093–35101 (2003).

35. Q. Zhu et al., Syntaxin-binding protein STXBP5 inhibits endothelial exocytosis and promotes platelet secretion. J Clin Invest 124, 4503–4516 (2014).

36. I. K. Madera-Salcedo et al., Tomosyn functions as a PKCδ-regulated fusion clamp in mast cell degranulation. Sci Signal 11, eaan4350 (2018).

37. M. J. Kennedy, I. G. Davison, C. G. Robinson, M. D. Ehlers, Syntaxin-4 defines a domain for activity-dependent exocytosis in dendritic spines. Cell 141, 524–535 (2010).

38. U. Ashery, N. Bielopolski, B. Barak, O. Yizhar, Friends and foes in synaptic transmission: the role of tomosyn in vesicle priming. Trends Neurosci 32, 275–282 (2009).

39. S. Ye et al., Platelet secretion and hemostasis require syntaxin-binding protein STXBP5. J Clin Invest 124, 4517–4528 (2014).

40. B. Barak et al., Tomosyn expression pattern in the mouse hippocampus suggests both presynaptic and postsynaptic functions. Front Neuroanat 4, 149 (2010).

41. A. J. Groffen, L. Jacobsen, D. Schut, M. Verhage, Two distinct genes drive expression of seven tomosyn isoforms in the mammalian brain, sharing a conserved structure with a unique variable domain. J Neurochem 92, 554–568 (2005).

42. J. Li et al., Spatiotemporal profile of postsynaptic interactomes integrates components of complex brain disorders. Nat Neurosci 20, 1150–1161 (2017).

43. B. M. Mechler, W. McGinnis, W. J. Gehring, Molecular cloning of lethal(2)giant larvae, a recessive oncogene of Drosophila melanogaster. The EMBO journal 4, 1551–1557 (1985).

44. K. Lehman, G. Rossi, J. E. Adamo, P. Brennwald, Yeast homologues of tomosyn and lethal giant larvae function in exocytosis and are associated with the plasma membrane SNARE, Sec9. J Cell Biol 146, 125–140 (1999).

45. M. Schapira, M. Tyers, M. Torrent, C. H. Arrowsmith, WD40 repeat domain proteins: a novel target class? Nat Rev Drug Discov 16, 773–786 (2017).

46. G. Rossi, K. Watson, M. Demonch, B. Temple, P. Brennwald, In vitro reconstitution of Rab GTPase-dependent vesicle clustering by the yeast lethal giant larvae/tomosyn homolog, Sro7. J Biol Chem 290, 612–624 (2015).

47. G. Rossi, K. Watson, W. Kennedy, P. Brennwald, The tomosyn homologue, Sro7, is a direct effector of the Rab GTPase, Sec4, in post-Golgi vesicle tethering. Mol Biol Cell 29, 1476–1486 (2018).

48. M. Kagami, A. Toh-e, Y. Matsui, Sro7p, a Saccharomyces cerevisiae counterpart of the tumor suppressor l(2)gl protein, is related to myosins in function. Genetics 149, 1717–1727 (1998).

49. V. A. Cazares et al., Dynamic Partitioning of Synaptic Vesicle Pools by the SNARE-Binding Protein Tomosyn. J Neurosci 36, 11208–11222 (2016).

50. N. R. Bin et al., Crucial Role of Postsynaptic Syntaxin 4 in Mediating Basal Neurotransmission and Synaptic Plasticity in Hippocampal CA1 Neurons. Cell Rep 23, 2955–2966 (2018).

51. Y. Gu et al., Differential vesicular sorting of AMPA and GABAA receptors. Proc Natl Acad Sci U S A 113, E922–931 (2016).

52. K. Hatsuzawa, T. Lang, D. Fasshauer, D. Bruns, R. Jahn, The R-SNARE Motif of Tomosyn Forms SNARE Core Complexes with Syntaxin 1 and SNAP-25 and Down-regulates Exocytosis. J Biol Chem 278, 31159–31166 (2003).

53. D. W. Allison, V. I. Gelfand, I. Spector, A. M. Craig, Role of Actin in Anchoring Postsynaptic Receptors in Cultured Hippocampal Neurons: Differential Attachment of NMDA versus AMPA Receptors. J Neurosci 18, 2423 (1998).

54. C.-H. Kim, J. E. Lisman, A Role of Actin Filament in Synaptic Transmission and Long-Term Potentiation. J Neurosci 19, 4314 (1999).

55. Q. Zhou, M. Xiao, R. A. Nicoll, Contribution of cytoskeleton to the internalization of AMPA receptors. Proc Natl Acad Sci U S A 98, 1261–1266 (2001).

56. E. Moretto, L. Murru, G. Martano, J. Sassone, M. Passafaro, Glutamatergic synapses in neurodevelopmental disorders. Prog Neuropsychopharmacol Biol Psychiatry 84, 328–342 (2018).

57. M. C. Subauste et al., Rho family proteins modulate rapid apoptosis induced by cytotoxic T lymphocytes and Fas. J Biol Chem 275, 9725–9733 (2000).

58. C. D. Kopec, B. Li, W. Wei, J. Boehm, R. Malinow, Glutamate receptor exocytosis and spine enlargement during chemically induced long-term potentiation. J Neurosci 26, 2000–2009 (2006).

59. T. Matsuda, C. L. Cepko, Electroporation and RNA interference in the rodent retina in vivo and in vitro. Proc Natl Acad Sci U S A 101, 16–22 (2004).

60. M. K. Bermejo, M. Milenkovic, A. Salahpour, A. J. Ramsey, Preparation of synaptic plasma membrane and postsynaptic density proteins using a discontinuous sucrose gradient. J Vis Exp 10.3791/51896, e51896 (2014).

61. E. Meijering et al., Design and validation of a tool for neurite tracing and analysis in fluorescence microscopy images. Cytometry A 58, 167–176 (2004).

62. T. A. Ferreira et al., Neuronal morphometry directly from bitmap images. Nat Methods 11, 982–984 (2014).

63. J. Noel et al., Surface Expression of AMPA Receptors in Hippocampal Neurons Is Regulated by an NSF-Dependent Mechanism. Neuron 23, 365–376 (1999).

